# Keras/TensorFlow in Drug Design for Immunity Disorders

**DOI:** 10.1101/2023.09.14.557712

**Authors:** Paulina Dragan, Kavita Joshi, Alessandro Atzei, Dorota Latek

## Abstract

Homeostasis of the host immune system is regulated by white blood cells with a variety of cell surface receptors for cytokines. Chemotactic cytokines (chemokines) activate their receptors to evoke the chemotaxis of immune cells in homeostatic migrations or inflammatory conditions towards inflamed tissue or pathogens. Dysregulation of the immune system leading to disorders such as allergies, autoimmune diseases, or cancer requires efficient, fast-acting drugs to minimize the long-term effects of chronic inflammation. Here, we performed structure-based virtual screening (SBVS) assisted by the Keras/TensorFlow neural network (NN) to find novel compound scaffolds acting on three chemokine receptors: CCR2, CCR3 and one CXC receptor CXCR3. Keras/TensorFlow NN was used here not as a typically used binary classifier, but as an efficient multi-class classifier that can discard not only inactive compounds but also low or medium-activity compounds. Several compounds proposed by SBVS and NN were tested in 100 ns all-atom molecular dynamics simulations to confirm their binding affinity. To improve the basic binding affinity of the compounds, new chemical modifications were proposed. The modified compounds were compared with known antagonists of these three chemokine receptors. Known CXCR3 were among the top predicted compounds and thus benefits of using Keras/TensorFlow in drug discovery have been shown in addition to structure-based approaches. Furthermore, we showed that Keras/TensorFlow NN can accurately predict the receptor subtype selectivity of compounds, for which SBVS often fails. We cross-tested chemokine receptor datasets retrieved from ChEMBL and curated datasets for cannabinoid receptors available at: http://db-gpcr-chem.uw.edu.pl. The NN model trained on the cannabinoid receptor datasets retrieved from ChEMBL was the most accurate in the receptor subtype selectivity prediction. Among NN models trained on the chemokine receptor datasets, the CXCR3 model showed the highest accuracy in differentiating the receptor subtype for a given compound dataset.

## 1. Introduction

Chemokines, or chemotactic cytokines, are a group of highly conserved small proteins that participate in the immune response via the chemotaxis of cells, either in response to tissue damage or infection (inflammatory chemokines) or to ensure homeostasis (homeostatic chemokines) [1], [2]. Inflammatory chemokines (incl. CCL1-13, CCL23, CCL24, CCL26, CXCL1-3, CXCL5-11) are expressed under inflammatory conditions and cause an increase in leukocyte trafficking towards the inflamed tissue. Homeostatic chemokines (incl. CCL14, CCL15, CCL16, CCL19, CCL21, CCL25, CCL27, CCL28, CXCL4, CXCL12, CXCL13), are expressed constitutively and induce constant homeostatic migrations to lymph nodes throughout the body and as well as the homing of immune cells such as lymphocytes [3]. However, dual-function chemokines, e.g., CCL11, CCL17, CCL22, or XCL1 (lymphotactin) and CXC3CL1 (fractalkine) also exist.

Due to the chemokine system being central to physiological processes such as homeostasis and immune responses by leukocyte transferring [4,5], the expression of these proteins a promising prognostic method for various malignancies [6]. Furthermore, a dysregulation of the chemokine system is implicated in cancer pathogenesis [7–9], but also in the progression of inflammatory and immune diseases, making chemokine receptors an emerging target in the development of new drugs.

The pro-inflammatory or homeostatic effects of chemokines are exerted through the activation of their receptors—a family of rhodopsin-like G protein-coupled receptors (GPCRs) that, based on the arrangement of cysteine residues in the N-terminal of the chemokines they bind, can be divided into four subfamilies: XCR, CCR, CXCR, and CX3CR [10]. To date, roughly 19 standard chemokine receptors and four atypical chemokine receptors (ACKRs) have been characterized in humans [11,12]. The latter group is less known and lacks a full-length structural characterization in the PDB but represents a promising group of drug targets. For example, ACKR3 modulates the CXCR4 signaling by acting as a decoy receptor and scavenging of CXCL12 [13]. The CXCL12 chemokine that binds both ACKR3 and CXCR4 is classified as a homeostatic chemokine and is over-expressed in autoimmune and inflammatory diseases [13]. Among homeostatic receptors, CXCR4, CCR7, and CCR9 are the most well-known [14] but many others are still being investigated.

CCR2 is a conventional chemokine receptor responding to chemokines with the cysteine CC motif in their N-termini. CCR2 is expressed largely in T cells and monocytes [15], and is specifically involved in monocyte mobilization [16]. Similarly to other chemokine receptors, CCR2 can be activated non-selectively by many different chemokines, including: CCL2, CCL7, CCL8, CCL12, CCL13, and CCL16 [16]. A recently discovered chemokine PSMP—PC3-secreted microprotein (microseminoprotein, prostate-associated MSMP), which is over-expressed in cancer and promotes hepatic fibrosis, has an affinity for CCR2 on a level similar to the most potent CCL2 [17]. This has implications for the importance of CCR2 in drug discovery for a variety of pathologies, e.g., inflammatory and autoimmune diseases such as rheumatoid arthritis [15], multiple sclerosis [16], and autoimmunity-driven type-1 diabetes [18], but also ischemic stroke [16], liver disease [19], asthma, atherosclerosis, transplant rejection [20], diabetic nephropathy, neuropathic pain, and the promotion of cancer cell metastasis [18]. CCR2 is the target of multiple clinical candidates—according to ChEMBL (accessed: July 2023) [21,22], nine are already in the 2^nd^ phase of clinical trials, and one more is in the 3^rd^ phase, but none has been approved for clinical use so far [23].

CCR3 belongs to the same subfamily of chemokine receptors as CCR2, and is expressed predominately on the surface of eosinophils [24] and basophils [25]. Although it is also over-expressed in certain types of cancer, it is connected with a rather poor prognosis (except in prostate and ovarian cancers) in contrast to a generally better prognosis associated with a high expression of CCR2 (except in glioma, testicular, and renal cancers) due to the CCR3-mediated migration of cancer cells [26]. CCR3 is known to bind chemokines CCL5, CCL7, CCL13, CCL15, eotaxin-1 (CCL11), eotaxin-2 (CCL24), and eotaxin-3 (CCL26) [25]. The activation of CCR3 by eotaxins (eosinophil chemotactic proteins) induces inflammation and thus is involved in asthma and allergies [24], including allergic skin diseases [27]. According to ChEMBL, there are currently no drugs nor clinical candidates targeting this receptor. Only recently have the active-state structures of CCR2 (bound to CCL2) and CCR3 (CCL11 not visible in electron density maps, but an active-like receptor structure) been solved using cryo-EM [23] to provide a basis for the rational design of novel immunomodulators acting on these two receptors.

CXCR3 is a chemokine receptor that is expressed mainly on immune cells such as natural killer cells and activated T lymphocytes. In humans, it can exist in three different isoforms: CXCR3-A, CXCR3-B, and CXCR3-alt, which are a result of gene splicing. While CXCR3-A and CXCR3-B sequences display a large overlap, there is a difference in their N-terminal, which is longer in the case of CXCR3-B due to the insertion of an additional sequence fragment from exon-2 of the CXCR3 gene. CXCR3-alt, however, consists of five transmembrane domains rather than seven as a result of the deletion of 337 base pairs from exon-3 [28]. Different isoforms are known to bind different chemokines—while CXCR3-A binds CXCL9, CXCL10, and CXCL11, CXCR3-B additionally binds CXCL4 [28]; CXCR3-alt is known to bind CXCL11 [29]. Similarly to CCR3, CXCR3 has been implicated in the progression of numerous diseases, including but not limited to multiple sclerosis, rheumatoid arthritis, transplant rejection [20], systemic lupus erythematosus [30], and allergies [31]. CXCR3 knockout mice are reported to be more resistant to autoimmune diseases [18]. A clinical candidate for acute lung inflammation targeting CXCR3 has been suggested by Meyer et al. but it has not yet been tested in clinical trials [32]. Biased ligands of CXCR3 (biaryl-type VUF10661 and VUF11418) have also been discovered in addition to the biased signaling observed for endogenous agonists of CXCR3 (CXCL11 bias towards β-arrestin). Recently, these three agonists of CXCR3 have been shown to activate the formation of the Gαi:β-arrestin complex in non-canonical GPCR signaling [33]. This emphasizes the importance of drug design for CXCR3 in numerous diseases, such as cancer, inflammatory diseases, and autoimmune disorders.

It is worth mentioning that except for the ACKRs, the structure of chemokine receptors is vastly conserved around the DRYLAIV motif in TM3 and the ICL2 loop [34,35]. However, only 3 drugs out of 45 in trials have so far been clinically approved [36]. Mogamulizumab was first approved in 2012 as a CCR4 antibody antagonist for cancer treatment. Maraviroc was approved in 2007 as an antiviral by acting as a CCR5 antagonist, while in 2008, Plerixafor was approved as a CXCR4 partial agonist for cancer therapies. To our knowledge, no drugs have been clinically approved so far for their action on CCR2, CCR3, or CXCR3.

As mentioned above, CCR2 is involved in a wide range of diseases, however, most of the clinical trials to find new CCR2-binding drugs failed in Phase II [37], [36]. CCR3 seems to be a target for asthma and allergy, but ongoing studies present a potential role of CCR3 antagonism in two disorders associated with the aging population, such as macular degeneration (MAD) and cognitive dysfunction in mice models [37]. CXCR3, mostly expressed on the surface of activated T cells, B cells, and natural killer cells, plays a crucial role in infection, autoimmune diseases, and tumor immunity by binding to specific receptors on target cell membranes to induce targeted cell migration and resulting immune responses. CXCR3 and its main ligands (i.e., CXCL9, CXCL10 and CXCL11) have been linked to the development of many tumors (Table 1). Interestingly, the CXCR3 ligands CXCL9, CXCL10, and CXCL11 demonstrate a dichotomous activity in cancer ranging from the inhibition to the promotion of tumor growth [38]. This can be explained by the varied expression patterns of CXCR3 in many tumor tissues. Therefore, it is necessary to better investigate the mode of action(s) (MoAs) and related signaling pathways for CXCR3 given its potential role as a new target for clinical tumor immunotherapy. Known antagonists and agonists of CCR2, CCR3, and CXCR3 receptors are shown in Table 1.

**Table 1.** Known active ligands of CCR2, CCR3 and CXCR3 receptors.

| Chemo-<br>kine re-<br>ceptor | Active ligands |  | Mechanism of Action | Indications | Status | Refer-<br>ences |
| --- | --- | --- | --- | --- | --- | --- |
|  | ligand | drugs |  |  |  |  |
| **<u>CCR2</u>** |  | CCX-140 | Antagonist | Type 2 diabetes and diabetic nephropathy | In trial, II clinical phase | [39]; [36] |
|  |  | Plozali-zumab (MLN-1202) | Antagonist | Anti-inflammatory | In trial, II clinical phase | [36] |
|  |  | Plozali-zumab, MLN-1202 | Antagonist | Antineoplastic | In trial, II clinical phase | [36] |
|  |  | CNTX-6970 | Antagonist | Analgesic | In trial, II clinical phase | [36] |
|  |  | Incb3284 | Antagonist | Anti-inflammatory | In trial, II clinical phase | [36] |
|  |  | azd2423 | Antagonist | Chronic obstructive pulmonary disorder | In trial, II clinical phase | [36] |
|  |  | Ccx872 | Antagonist |  | In trial, II clinical phase | [36] |
|  |  | cenicriviroc | Antagonist | Antiviral, HIV | In trial, II clinical phase | [36] |
|  |  | Ccl2-lpm | Antagonist | Anti-inflammatory | In trial, II clinical phase | [36] |
| **<u>CCR3</u>** |  | Tpi-asm8 | Antagonist | Anti-asthmatic | In trial | [36] |
| **<u>CXCR3</u>** | CXCL9/10 |  | Promoting lymph node metastasis; Promotion of lymph node and lung metastasis; Promotion of malignant ascites production; Promoting tumor growth and metastasis; respectively | Colorectal cancer<br>Breast cancer; Ovarian cancer; Lung cancer; Stomach cancer; |  | [40] |
|  | CXCL9/10/11 |  | Promotes proliferation and metastasis of cancer cells; Promotes distant metastasis; Inhibit tumor growth and metastasis; respectively | Esophageal cancer; Kidney cancer; Osteosarcoma |  | [40] |
|  | CXCL10 |  | Promotes tumor growth and metastasis; Inhibits tumor growth; respectively | Prostate cancer; Glioma and Myeloma |  | [40] |

Drug discovery is a long and costly process, but it can be enhanced by computational methods, including both structure- and ligand-based virtual screening (SBVS and LBVS, respectively). Of these two, SBVS is more time-consuming, requiring the use of a target structure or a homology model to compute the approximate free energy of the ligand binding [41]. The aim of SBVS is to screen a library of compounds using a receptor homology model or its cryo-EM/X-ray structure using scoring functions based on simplified force fields. The computed free energy of binding, approximated by a scoring function (SF), enables the selection of compounds that will likely evoke the highest response *in vitro*. Many different programs are available to perform such library screening, including Glide [42], AutoDock, AutoDock Vina [43,44], DOCK [45], MOE [46], GOLD [47]. Comparative studies done on AutoDock and AutoDock Vina indicate that the latter is better at predicting binding poses, though it cannot be said that one program is inherently superior to the other—they were found to be better fitted for different drug targets. The computational time is also crucial to deciding which docking program to choose for virtual screening purposes. Recently, Autodock Vina has been modified to fit the GPU architecture, which significantly accelerates the computations and adds more advantages in comparison to other molecular docking software [48].

In recent years, scoring functions (SFs) used in SBVS have also begun to be based on machine learning. In contrast to classical SFs, ML-based SFs do not make use of a fixed functional form (usually linear) that is based on the relationship between the characteristics of the protein-ligand complex and the binding affinity. Instead, in their case the functional form is based purely on the information obtained by ML from the training data [49]. This way, it is possible to reflect non-linear relationships between the protein-ligand complex structure and the ligand binding affinity, e.g., by using neural networks (NNs), random forest (RF), or support vector machines (SVM) [50]. Deep learning methods—especially convolutional neural networks (CNN)—have been applied in SBVS in order to obtain more reliable results from docking calculations [50]. Such approaches include DeepVS [51], DenseFS [52], and Gonczarek et al.’s fingerprinting method involving learnable atomic convolution [53]. Furthermore, CNNs have also been used in the prediction of binding poses and affinities, and more robust models can be built by combining them with transfer and multitask learning [50]. Noteworthily, deep learning methods are not always better than those based on classical machine learning [54]. Classical ML methods are typically used for rescoring or ranking of the output from popular molecular docking programs rather than being directly integrated into them, and their results are not easily interpretable [50]. The interpretability of results of a deep learning method can be important especially when it comes to medical applications [55], e.g., for finding gene-drug associations [56]. One of the common methods to explain ML results is SHapley Additive exPlanations (SHAP) [57]. Shapley values allow the importance of specific features to be assessed by computing three properties: consistency, missingness, and local accuracy. This method demonstrates a high consistency with human intuition [57]. Other interpretability methods include DeepLIFT [58], especially used for deep NNs, or Grad-CAM++ [59], used to visually explain the predictions of CNN models [57]. There are also frameworks joining various methods to uncover global feature importance in contrast to local interpretation of each feature, e.g., SAGE [60,61].

In principle, LBVS is much faster, based solely on the structure and physicochemical properties of ligands known to interact with the molecular target in order to predict the affinities of yet untested compounds [62]. This makes it possible to use when the structure of a receptor is unavailable, which is often the case with GPCRs. The applicability of machine learning methods in ligand-based virtual screening has been widely discussed so far, e.g., in [63]. Constantly increasing in the number of available ligand datasets for various drug targets and improving of the quality and quantity of such datasets improves the accuracy of computational drug discovery despite minor problems with integration and optimization of used ML methods [64]. In supervised ML, feature selection is used to recognize relevant molecular (in case of drug discovery) or genomic (in case of genomic analysis) features that inform about drug responses or drug-gene associations. These techniques, however, require the labeling of used training dataset, e.g., prior knowledge about drug-target associations [65]. Thus, their use may inhibit the discovery of potential new actives, as compounds not possessing the preselected features could be discarded.

Data-driven concepts to result in, e.g., new drugs or drug targets, require efficient and accurate algorithms in advance, to process massive data from large biomedical repositories and to reflect subtle differences in compounds that have a huge impact on the observed biological response, respectively. Although conventional ML algorithms belonging to a supervised learners’ group, such as gradient boosting or support vector machines, seem to be the most accurate in tasks including predictions of compound activity or binding affinity [54,63,66–68], NNs constantly draw attention [64]. Learning of NNs can be carried out as supervised learning (first NNs, also including backpropagation NNs), or unsupervised learning (e.g., deep belief networks) where layers can detect relevant features. Other types of learning, e.g., reinforcement learning can also be used to train NNs [69]. In principle, if no labeled input data is used to train NN, it has many more possibilities of finding new active-like scaffolds compared to supervised-learning methods. This is the basis for the popularity of NNs and deep learning NNs in various tasks, ranging from image recognition, natural language processing, and engineering applications [70], to the retrieval of relevant information from databases, e.g., in protein structure prediction [71] or in drug design [72]. The two, mostly used systems for machine learning and especially for deep learning tasks are TensorFlow, developed by Google [73], and PyTorch [74], co-developed by A. Paszke. Keras API [75] makes it possible to define and train ML models implemented on TensorFlow or PyTorch platforms to easily release open-source projects and construct pipelines joining various libraries, e.g., RDKit [76] for compound fingerprints [77]. TensorFlow with or without high-level Keras API is widely used due to the easy implementation of algorithms that are otherwise difficult to optimize flawlessly, such as for example convolutional neural networks [78]. One of the key concepts recently introduced in TensorFlow2.0 and PyTorch is a ‘define-by-run’ paradigm [79,80], in which connections in NNs are defined during the training, not before. This backpropagation allows for a more efficient automatic differentiation scheme compared to ‘define-and-run’ in TensorFlow1.0.

Recently, TensorFlow has been used for rapid screening for GPCR ligands [81] but the combination of two methods, LBVS and SBVS, allows for a more precise and reliable assessment of the ligand binding affinity and its detailed binding mode [54]. In the final step, molecular dynamics (MD) can be used to validate the molecular docking-based binding affinity and binding modes of discovered compounds and thus to reduce the number of false positives before the bioassay studies [82]. This combined computational approach significantly reduces both time and cost required to find novel chemotypes.

Here, we performed MD simulations for previously obtained novel CCR2 and CCR3 antagonists, and used a combination of AutoDock Vina for SBVS, Keras/Tensorflow sequential model of neural network (NN) for LBVS, and MD simulations in order in order to find and validate novel small-molecule antagonists for CCR2, CCR3, and CXCR3 chemokine receptors. While previously [81] the impact on the ligand dataset composition on the ML results was discussed, here we focused on the ability of ML to reflect slight structural differences between ligands matching the certain receptor subtype which account for their receptor subtype selectivity. In [83] we assessed gradient boosting decision trees (LightGBM) in the recognition of the receptor subtype selective and non-selective ligands of cannabinoid receptors. Here, we assess NNs (Keras/TensorFlow) in such a task using not only the curated ligand datasets for CB1/CB2 cannabinoid receptors (http://db-gpcr-chem.uw.edu.pl), but also for CCR2/CCR3/CXCR3 chemokine receptors.

## 2. Results

### 2.1. CXCR3 Model Validation

The positions of transmembrane helices 5 and 6 (TM5 and TM6) were the first to be analyzed. It is well-known that during the activation of the GPCR receptor, the extracellular region of its TM5 helix moves inwards, while the intracellular region of TM6 moves outwards [84]. These differences in the inactive and active-state conformations are shown in Fig. 1. Indeed, the location and shape of TM5 and TM6 in our CXCR3 model are similar to those observed for inactive-state structures of chemokine receptors.

**Figure. 1.**
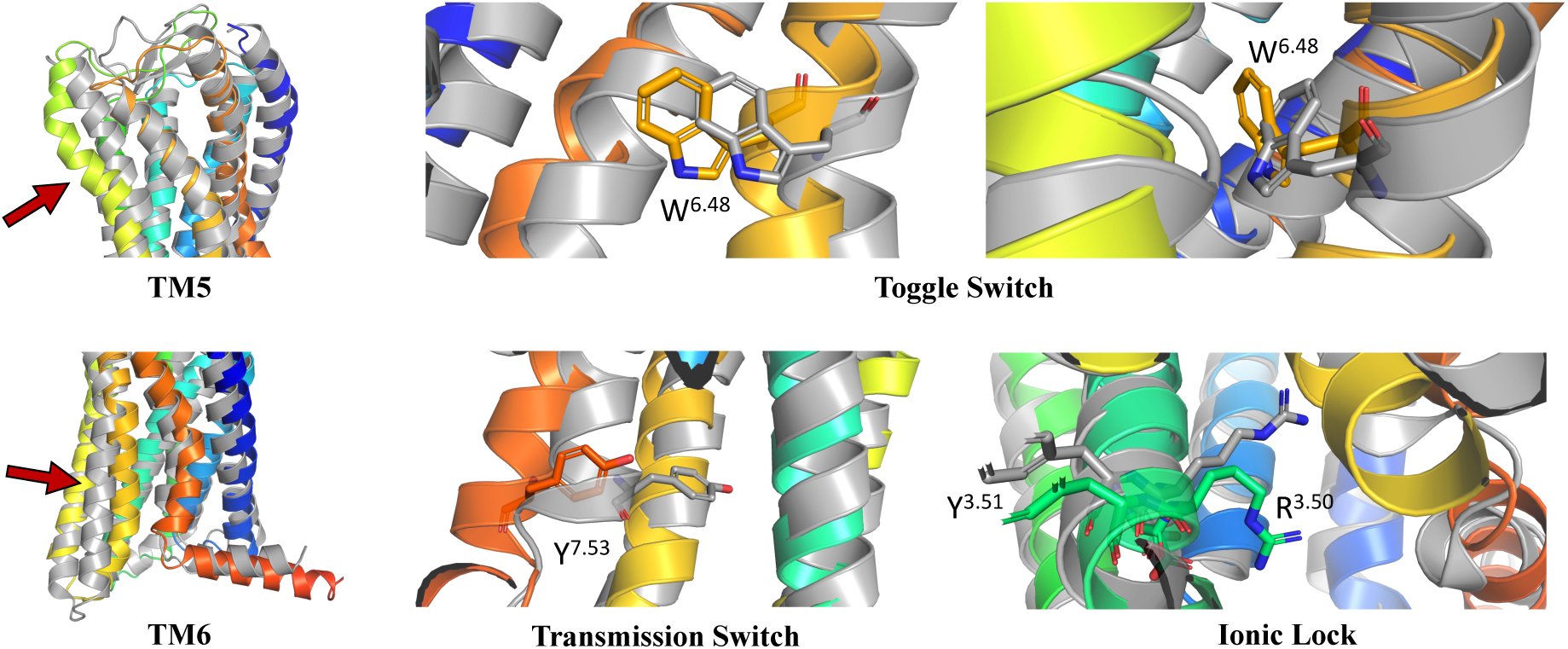
The validation of the inactive-state CXCR3 model through an analysis of micro- and macroswitches. The inactive-state model of CXCR3 (blue-to-red) was superposed on active-state chemokine receptor structures (gray): 7O7F (CCR5) for TM5 and TM6, and 6WWZ (CCR6) for comparison of TM helices and microswitches (toggle and transmission switches, and the ionic lock). The residues have been labeled using the Ballesteros-Weinstein numbering system [87].

Similarly, the conformations of selected residues were analyzed: the W^6.48^ toggle switch, Y^7.53^ transmission switch, and D^3.49^R^3.50^Y^3.51^ ionic lock [84–86]. The W^6.48^ toggle switch was rotated as compared to the active-state structure, the Y^7.53^ transmission switch had an altered position, and R^3.50^ in the ionic lock was bent inwards rather than straightened out. On this basis, it was concluded that the model was suitable for performing structure-based virtual screening, i.e., the molecular switches present in the model were in the conformations expected to be present in an inactive-state chemokine receptor.

### 2.2. MD-based validation of ligand binding modes

CCR2 and CCR3 actives proposed in a recent study by Dragan et al. were docked to the same receptor structures as before but with a different algorithm (Autodock Vina) to confirm their binding modes. Based on these results, 6 out of 10 CCR2 actives, and 7 out of 12 CCR3 actives were discarded as Glide and Autodock Vina provided significantly different binding modes for them. For CXCR3 only Autodock Vina was used for molecular docking in prior to MD simulations. Notably, molecular docking algorithms, extremely useful for virtual screening, have limitations regarding their reproducibility of ligand binding modes. This is due to simplified force fields, in which some of molecular interactions are approximated to decrease the computational time and to efficiently screen large libraries of compounds [63]. For this reason, all-atom MD simulations were used to validate the binding modes of proposed active compounds (see Appendix S1 Table S1-S3), following a previous study [88,89]. Of the four ligands tested for CCR2, only three remained stable throughout the course of the simulation (Fig. 2). Both Z144527132 and Z199951150 displayed a high stability from the very beginning of the production run— both the RMSD values and their standard deviations were low. Z2607653068, on the other hand, was stable at the beginning, but its hydroxyl group began moving upwards after 60 ns, as though to leave the receptor. At the 100 ns cutoff, the only interaction noted by Maestro for this ligand was pi-pi stacking with Y^3.32^; at this point in the simulation, the other tested molecules all displayed more binding interactions, further undermining the Z2607653068 binding mode obtained from molecular docking. The final compound, Z45637008, stabilized after 10 ns of the simulation, but after a noticeable relocation of its trifluoromethyl end. This change made it possible for the sulfonyl group and a nearby nitrogen atom to form hydrogen bonds with C^5.26^. Overall, pi-pi stacking interactions with W^2.60^ and Y^3.32^ appeared in multiple cases, suggesting these residues may play a significant role in the binding of CCR2 ligands.

**Figure. 2.**
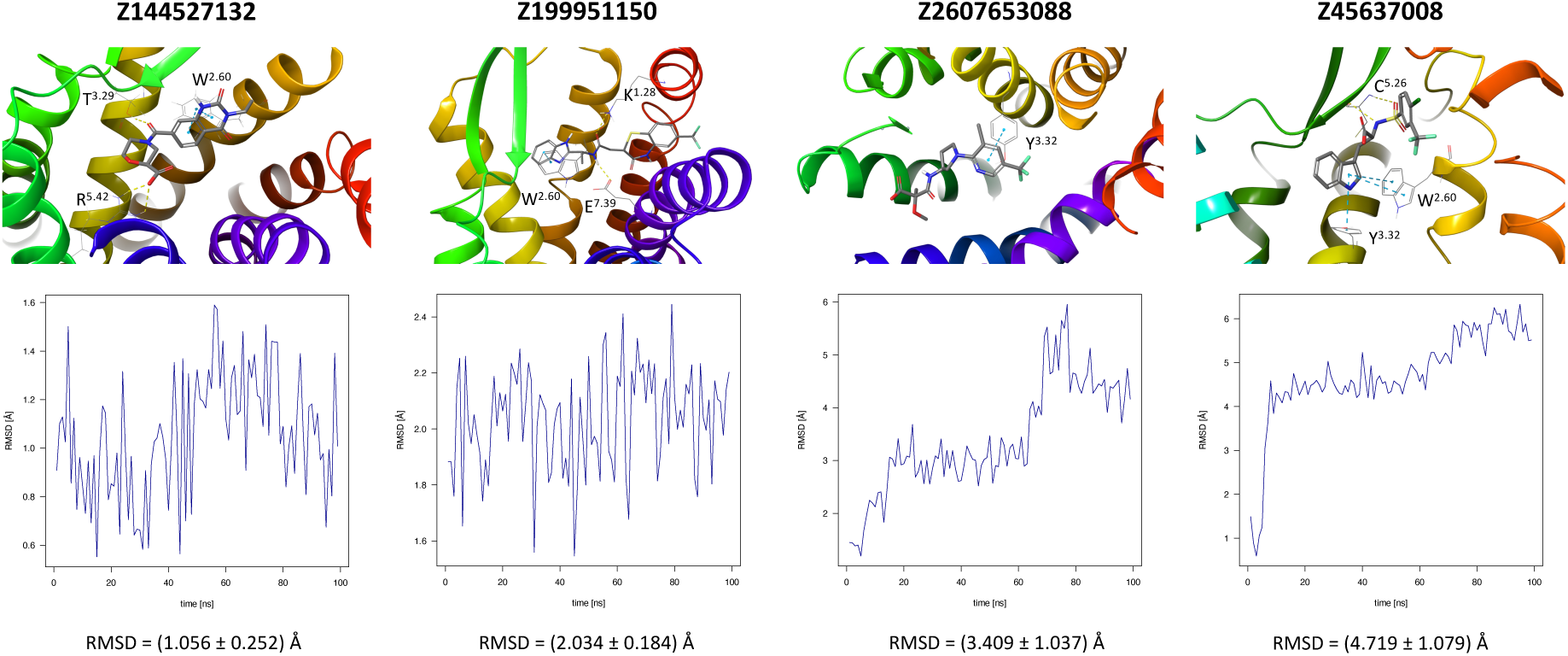
Results of the MD simulations for CCR2 for four different compounds proposed by virtual screening. Top: the interactions between the receptor and the ligand obtained after 100 ns of the simulation. The receptor was shown in the red-to-blue color scheme; yellow dashed lines—hydrogen bonds; blue dashed lines—pi-pi stacking. The residues have been labeled using Ballesteros-Weinstein numbering system [87]. Bottom: the RMSD plots obtained for each of the ligands over the 100 ns simulation, as well as the average RMSD with its fluctuation range.

In the same way, five molecules were tested for the inactive-state CCR3 Robetta model (Fig. 3). Z1912507172 was highly unstable over the first ca. 40 ns of the simulation. The ligand was observed to move much deeper into the binding site than according to the molecular docking results. After 40 ns its location stabilized and remained that way until the end of the simulation. A comparison of its binding mode, both obtained in molecular docking and refined in the MD simulations, are presented in Appendix S1 Table S2. Of the tested CCR3 compounds, Z2441027668 was the most stable, barely changing its position with respect to the results from molecular docking. Slightly larger, though still low, RMSD fluctuations were observed for the remaining three compounds: Z2764968046, Z1274732994, and Z2606182917. In general, the RMSD fluctuations obtained for the CCR3 complexes were higher than those obtained for CCR2 complexes, likely because a receptor model was used here rather than a high-quality structure. Similarly, as in the case of CCR2, W^2.60^ and Y^3.32^ were shown to frequently participate in ligand binding; in addition, interactions with E^4.60^ were noted in two separate cases.

**Figure. 3.**
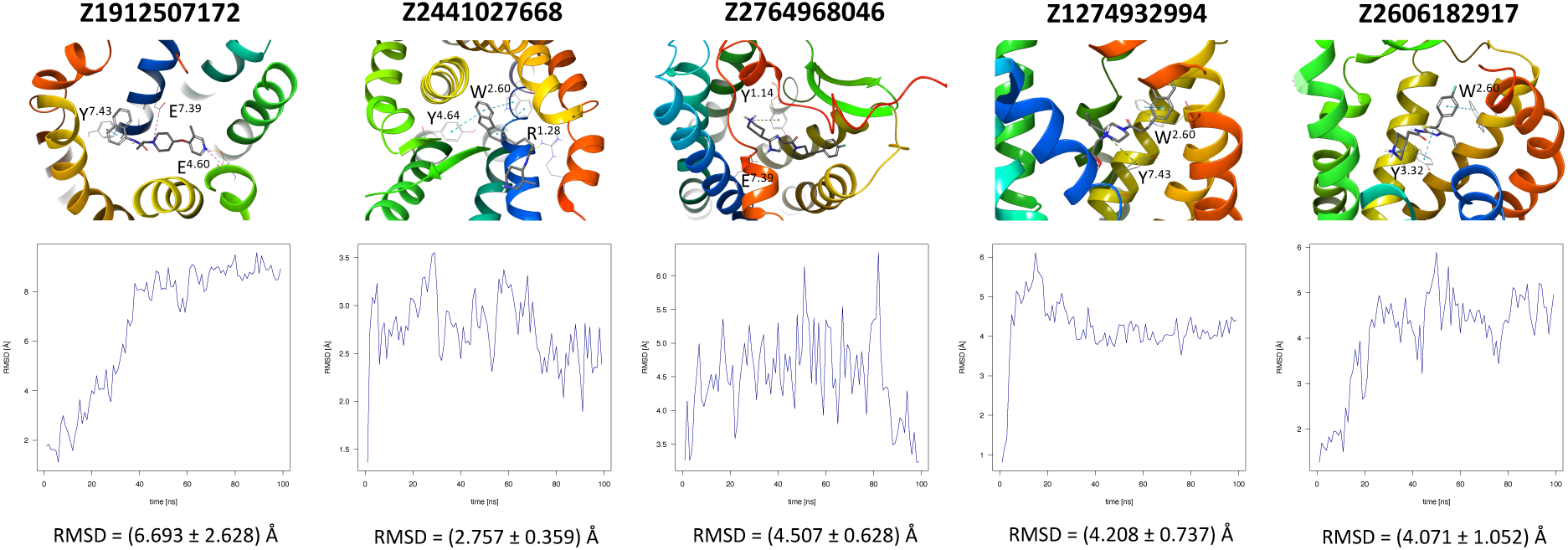
Results of the MD simulations for CCR3 for five proposed compounds. Top: the interactions between the receptor and the ligand obtained after 100 ns of the simulation. The receptor was shown in the red-to-blue color scheme; yellow dashed lines—hydrogen bonds; blue dashed lines— pi-pi stacking; green dashed lines—pi-cation; purple dashed lines—salt bridges. The residues have been labeled using Ballesteros-Weinstein numbering system [87]. Bottom: the RMSD plots obtained for each of the ligands over the 100 ns simulation, as well as the average RMSD with its fluctuation range.

A further five compounds were tested for the CXCR3 model—they all demonstrated a high stability of their binding mode. The most interactions with the receptor were observed for Z2233592864—mainly pi-pi stacking. This compound, along with Z107207944 and Z1903257002, demonstrated the most stable binding mode. The two final compounds, Z1167188972 and Z1510954688, fluctuated to a much greater extent, and no specific interactions with the receptor were observed for the latter one, while for the former interactions with residues N^3.33^ and Y^3.37^ were rarely formed during the simulations.

Interestingly, the best compounds selected by the Keras/TensorFlow NN for CXCR3 (in the range of 9 and above of pChEMBL predicted values) did not include any of the compounds proposed for CCR2 or CCR3 (Fig. 2 and 3). It means that like previously [54], predictions made by the NN model are selective for the receptor subtype because they are based on known active ligands only and not on the receptor structures which could be too similar, e.g., for SBVS. Thus, diverse CCR2, CCR3, or CXCR3 ligand training sets will provide diverse novel chemotypes for each of these receptors, while the similar structures of these receptors could only provide similar compounds in SBVS. This is further discussed in 2.5.

### 2.3. Chemical Modifications of Proposed Ligands

Chemical modifications of the structures of the proposed compounds were suggested in order to increase their affinity towards the receptors. The modified structures as well as their contacts with the receptor were presented in Appendix S1 Table S4-S6. In the case of the proposed CCR2 ligand Z144527132, the addition of the alkyl substituent with the hydroxyl group to the hydrogenated quinazoline ring was proposed in order to enable the formation of a hydrogen bond with E^7.39^, a residue that participated in the binding of another ligand for this receptor. A second alkyl group was added in order to fill out the hydrophobic pocket. For Z199951150, a shortening of the molecule (specifically, of the end with the trifluoride group) could be suggested, as it did not appear to greatly contribute to the binding. More significant modifications should be introduced to the structure of Z2607653068 in order to prevent it from immediately leaving the receptor. A phenyl ring in the place of the ligand part with carbonyl and hydroxyl groups might support the formation of pi-pi stacking interactions with the nearby F5.30 or Y^5.29^. Furthermore, an introduction of a cyclopentene ring between one of the carbons and an oxygen could facilitate the formation of a hydrogen bond with H^5.38^. No modifications were suggested for Z45637008. However, only one modified compound based on Z2607653068 (together with Z199951150) was in the 8-9 predicted activity range by Keras/TensorFlow NN, while the other two (see Appendix S1 Table S4) were predicted as inactive (below 5).

For CCR3, the introduction of a hydroxyl group into the structure of Z1274732994 was suggested to facilitate the formation of a hydrogen bond with E7.39. In the case of Z1912507172, the addition of two separate alkyl groups were suggested in order to better fill out the hydrophobic region of the binding pocket of the receptor, as well as a hydroxyl group that could interact with Y^3.32^ to form a hydrogen bond. For Z2441027668, it was suggested that the methylpiperidine ring could be transformed into methylpyridine in order to allow for potential pi-pi stacking interactions with Y^1.14^. For Z2606182917, the addition of an alkyl chain is suggested in order to better fill the binding cavity, as well as an oxygen that could form a hydrogen bond with H^5.38^. In the case of Z2764968046, a cyclopentane ring was added to the structure in order to better fill out the binding pocket. All modified compounds were in the highest predicted activity range (above 9 or in the 8-9 range) except for Z2606182917 that fell into the medium predicted activity range (7-8).

For CXCR3, an additional double bond to introduce aromaticity was added to the indane ring of Z107207944. Thus, the formation of pi-pi stacking interactions with the nearby F^3.32^, W^6.48^, or Y^6.51^ could be facilitated. For Z1167188972, a benzene ring could be a replacement for the cyclohexane ring to facilitate the pi-pi stacking interactions with F^4.63^. Furthermore, an alkene chain was added to fill out the binding pocket. In the case of Z1510954688, the tetrahydropyran ring can be replaced with a benzene ring, and one of the methyl groups was removed. This would allow for pi-pi stacking interactions with Y^3.37^. In addition, a transformation of one of the other methyl groups present in the molecule into a hydroxyethyl group would allow for the formation of a hydrogen bond with D^4.60^. For Z1903257002, the cyclohexane ring could be replaced with a benzene ring to facilitate interactions with Y^6.51^. A subsequent relocation of one of the methyl groups would help fill out the binding pocket. No modifications were suggested for Z2233592864. Interestingly, this compound (Z2233592864) together with a modified Z107207944 were the best among all modified compounds according to Keras/TensorFlow NN (the 8-9 predicted activity range).

### 2.4. Comparison of proposed compounds with known CXCR3 ligands

Recently, Meyer et al. [32] published a novel CXCR3 antagonist. A comparison of the described structures and Enamine’s Hit Locator Library (HLL) provided three hits. Two of the molecules, Z2755039307 and Z2755039304 proved most similar to ACT-7779991 (the clinical candidate) with Tanimoto similarities equal to 0.255 and 0.253, respectively, as well as ACT-672125, with similarities equal to 0.141 for both. A third compound, Z1695828968, was most similar to ACT-660602, with a similarity of 0.207. Here, the previous two compounds had similarities equal to 0.196 and 0.195, respectively. Autodock Vina-approximated binding affinities for these similar compounds in Enamine HLL were rather low to medium, ranging from 6.5 to 7.5. For two compounds, Z2755039307 and Z2755039304, the NN results were also unsatisfactory (see Table 2). However, the third compound Z1695828968 was assessed by the Keras/TensorFlow NN as highly active (the activity range above 9—the highest one) and it was included in less than 20 % of the best compounds for CXCR3 in Enamine HLL. It shows that the Keras/TensorFlow NN does not reproduce molecular docking results but indeed may provide substantial new information on the compound activity not accessible to physics-based force fields. To compare, the NN results for CXCR3 antagonists proposed and tested in MD simulations in this study fell into 18.6%, 3.7%, 13.5%, 19.5%, and 14.0% of top NN predictions (see Fig. 4, respectively) and 72.5%, 14,4%, 52.6%, 76.1%, and 54.9% of top predictions of the 9 and above activity range, respectively.

**Table 2.**
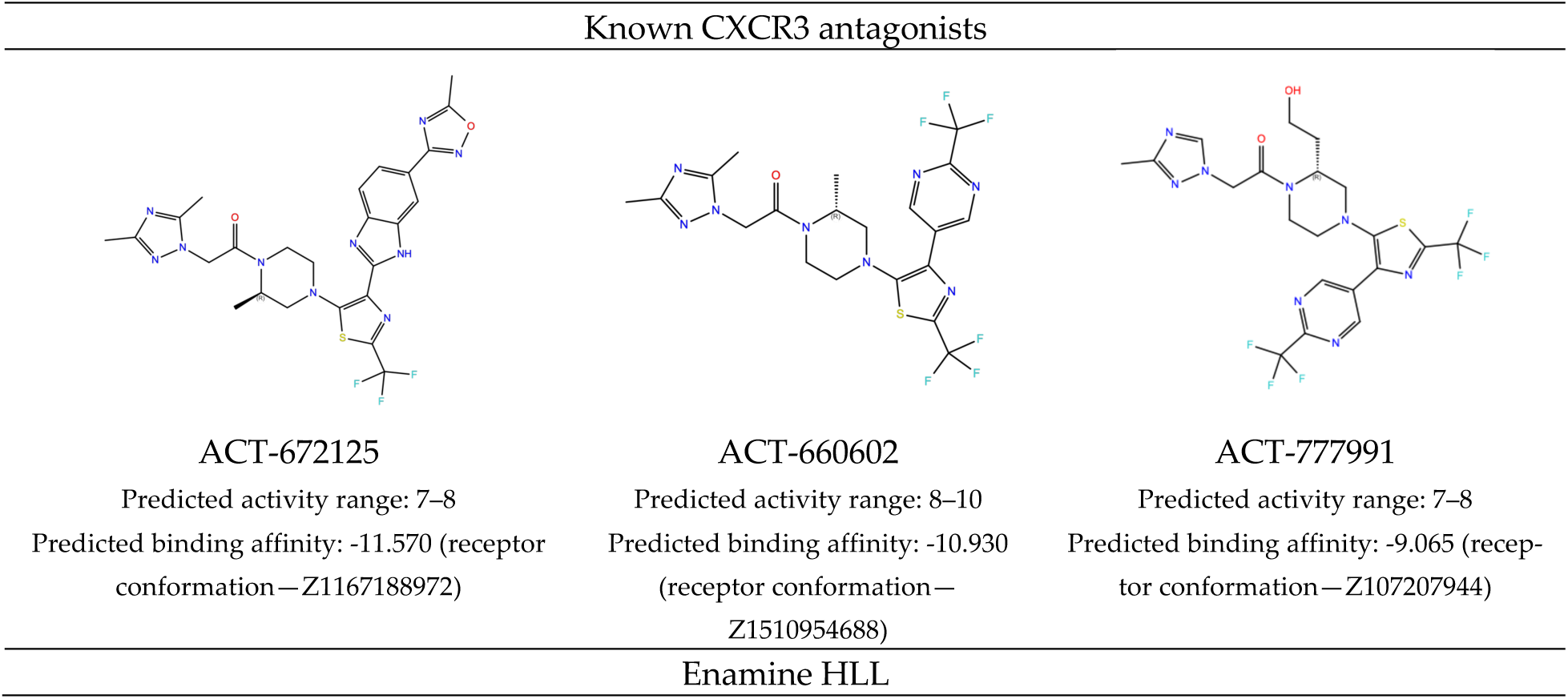
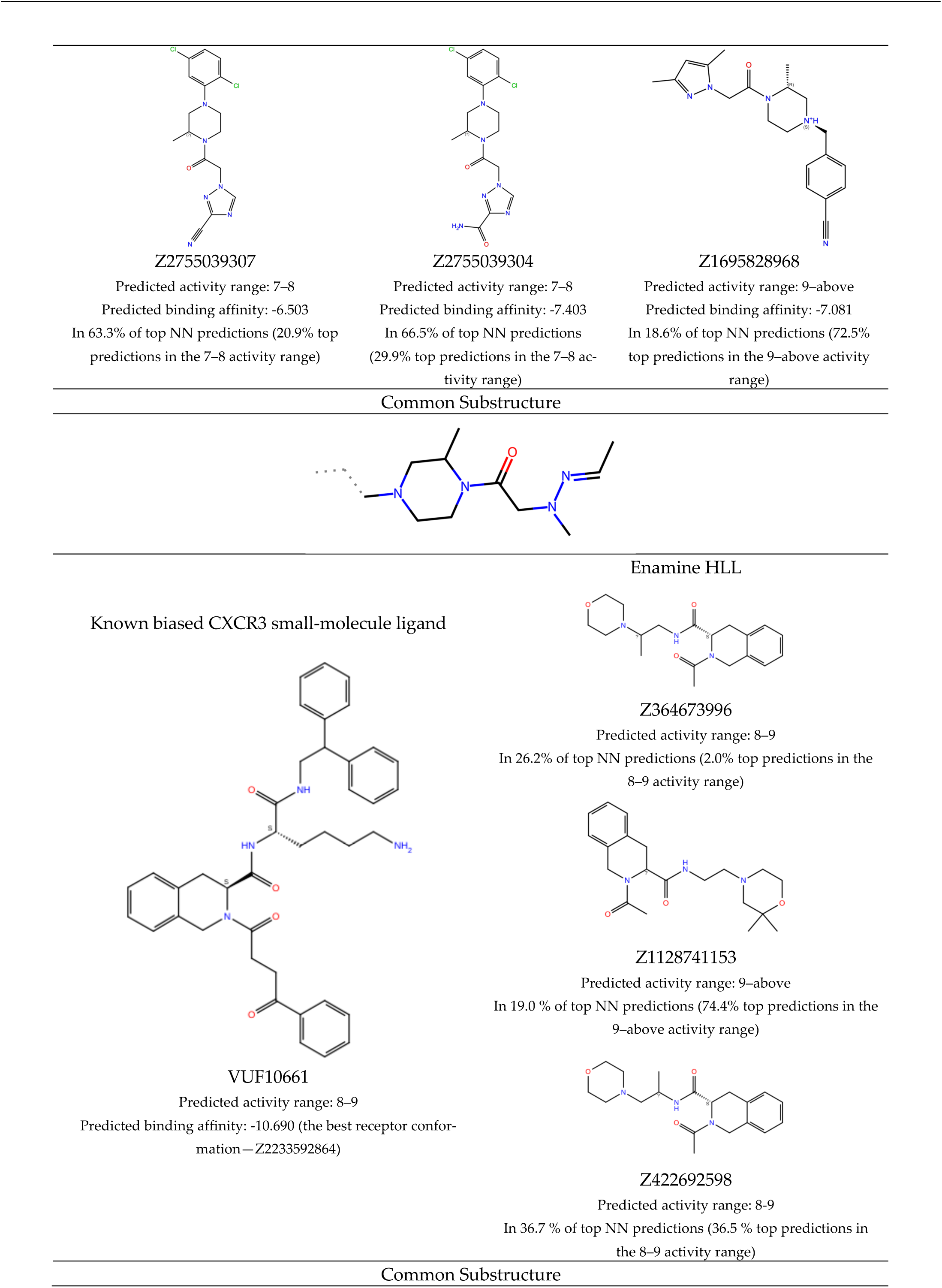
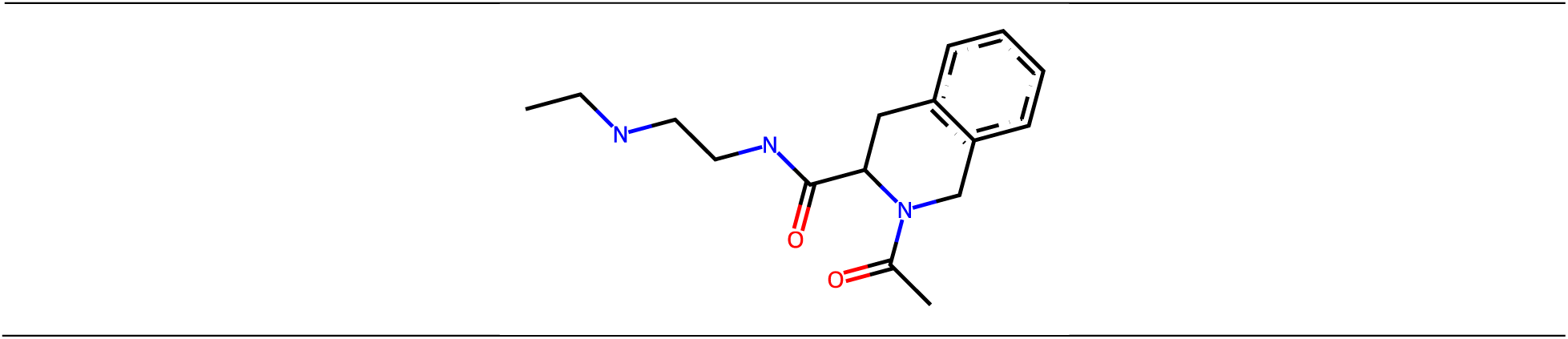
A comparison of three known CXCR3 active compounds proposed by Meyer et al. and a biased CXCR3 ligand with the most similar compounds present in Enamine’s Hit Locator Library. The common substructure of all six compounds is presented in the last row.

**Figure. 4.**
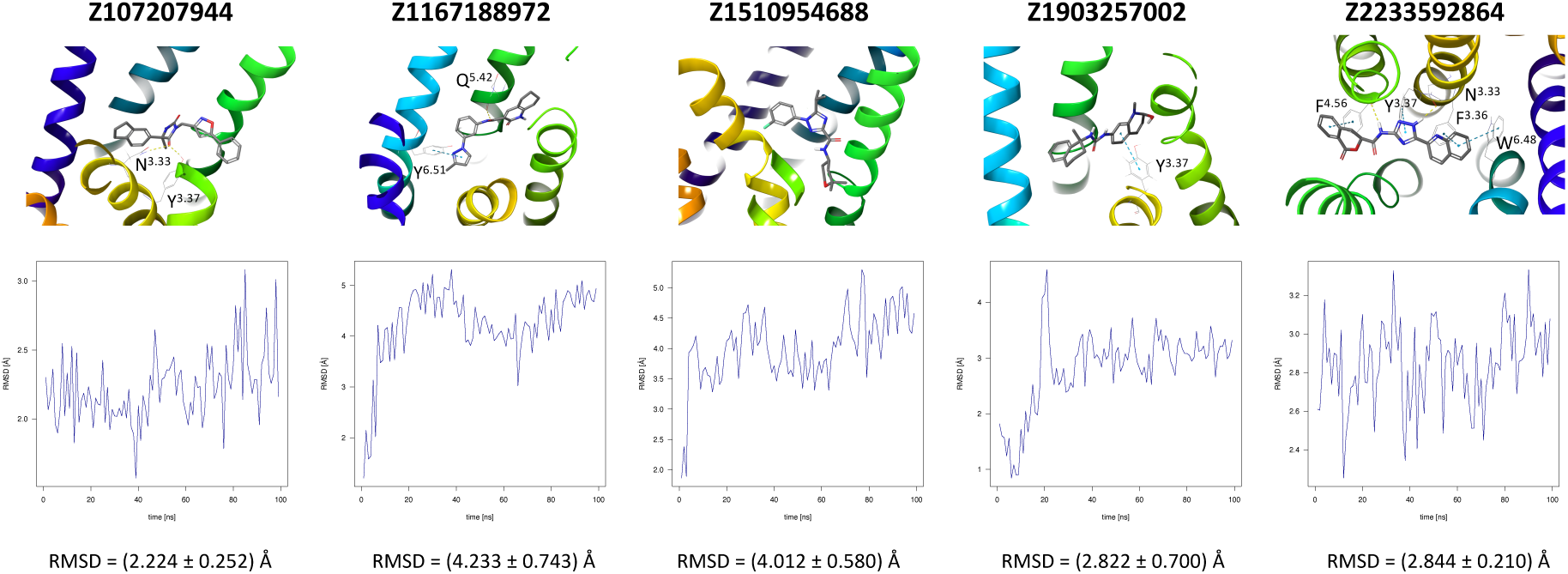
Results of the MD simulations for CXCR3 for five proposed compounds. Top: the interactions between the receptor and the ligand obtained for 100 ns of the simulation. The receptor was shown in the red-to-blue color scheme; yellow dashed lines—hydrogen bonds; blue dashed lines— pi-pi stacking. The residues have been labeled using Ballesteros-Weinstein numbering system [87]. Bottom: the RMSD computed for each of the ligands over the 100 ns simulation, as well as the average RMSD with its fluctuation range.

In addition to the above CXCR3 antagonists, we also searched for the compounds similar to a CXCR3 biased ligand VUF10661. Here, the results of NN were even better (see Table 2). All three similar compounds were in the activity range of predicted pChEMBL values of 8 and above, meaning they were predicted as highly active for CXCR3. All three of these compounds were also among the best compounds found in Enamine HLL. Despite these results, in our opinion, Keras/TensorFlow NN is a method to be used in combination with classical virtual screening methods such as SBVS rather than to be used solely in VS.

All four known CXCR3 ligands were docked with AutoDock Vina to five receptor conformations obtained at the end of MD simulations of five HLL compounds (see Fig. 4). The best binding affinities predicted by AutoDock Vina are shown in Table 2, while the binding modes are shown in Appendix S1 Table S7. There was no common receptor conformation that proved the best fit for all four compounds, but the Z1903257002-fitted conformation was discarded by all compounds possibly because of the steric hindrance caused by Y^3.37^ forming interactions with Z1903257002 (see Fig. 4).

All four CXCR3 ligands were additionally assessed by NN trained on the CXCR3 dataset and the CCR2 and CCR3 datasets to check if NN is sensitive to the receptor subtype selectivity. The CXCR3 model assessed these ligands rather highly, similarly as the CCR3 model, while the CCR2 model assessed them as inactive compounds. A similar observation was made for the modified compounds. They were among the top-scored compounds assessed by the CXCR3 and CCR3 models, but not by the CCR2 model. This means, that the CCR3 and CXCR3 NN models were less selective with respective to each other in the activity predictions for these compounds in comparison to the CCR2 model. This again suggests the dependency of NN on the training dataset composition [54], yet in this case with the desired outcome.

The NN and SBVS predictions were not fully consistent for Meyer’s compounds, meaning that the best compound proposed by NN was not the best compound proposed by SBVS. However, both NN and SBVS assessed VUF10661 as the best compound out of these four actives. This could be due to the fact, that VUF10661 consists of much more functional groups than Meyer’s compounds. More functional groups decrease the energy of interactions computed in molecular docking as observed previously by us in the statins case [90]. On the other hand, the presence of more functional groups ensures that the compound resembles at least any subset of active compounds used for training of NN and thus NN will select it as an active compound.

### 2.5. Performance of Keras/TensorFlow NN in the receptor subtype selectivity prediction tasks

To compare with the previous ML study on cannabinoid receptors (LightGBM, CB1/CB2 selectivity) we also used CB1 and CB2 datasets for the NN training. This time, we included as many ChEMBL-retrieved compounds as possible (> 5000) in contrast to previous limited datasets for these two receptors [83] available at: https://db-gpcr.chem.uw.edu.pl. The average Tanimoto coefficients between the current datasets and the previous datasets were equal to 0.138 (mode: 0.17) and 0.141 (mode: 0.17) for CB1 and CB2, respectively. Both datasets included small-molecule compounds only. However, the previous datasets included data from assays that provided pKi values, while the current datasets included only data from assays that provided pIC50 (standardized to pChEMBL values). This means that the current datasets include only CB1 or CB2 small-molecule inhibitors and not all CB1 and CB2 actives like previousely. Furthermore, the previous datasets did not include any inactive or weakly active compounds (pChEMBL < 4), the addition of which to training sets was recently discussed in [54]. In the current datasets, nearly 40 % and 25 % (CB1, CB2, respectively) of compounds were inactive compounds (pChEMBL equal to 0). Among active compounds in the current datasets 1 % and 2 % were weakly active compounds (pChEMBL less than 5, CB1 and CB2, respectively). Histograms showing distribution of the activity classes in the current and previous datasets were shown in Appendix S1 Table S8. Despite these differences, the results of the receptor subtype selectivity prediction tasks were similar for the current and previous datasets, with only a slight improvement in comparison to the previous ones. The accuracy of the prediction for validation datasets were ca. 0.5 for the same receptor subtype, 0.2 for the other receptor subtype, less than 0.02 for CB2 selective compounds with the inconsistent receptor subtype and near 1 for the consistent receptor subtype (see Table 3 and 4). In the latter case of the CB2 selective compounds with the matching receptor subtype, the CB2 model trained on the previous dataset performed much better than trained on the current dataset (accuracy: 0.876 vs. 0.521, see Table 4). This is however could be due to the higher average similarity between the previous training sets and the CB2 selective set (0.153 and 0.141 for the previous training set and the current one, respectively, see Table 4).

**Table 3.** Performance of Keras/TensorFlow NN in the chemokine receptor subtype selectivity tasks.

| Training set | Number of data-points | Validation set | Number of data-points | Loss | Accuracy (change) | Average Tanimoto coefficient<br>training vs. validation set | Mode Tanimoto coefficient<br>training vs. validation set |
| --- | --- | --- | --- | --- | --- | --- | --- |
| CCR2 | 1995 | CCR2 | 399 | 5.406 | 0.190 | 0.139 | 0.16 |
|  |  | CCR3 | 121 | 9.621 | 0.231 (+0.041) | 0.141 | 0.16 |
|  |  | CXCR3 | 199 | 17.878 | 0.126 (-0.064) | 0.125 | 0.13 |
| CCR3 | 603 | CCR3 | 121 | 5.332 | 0.223 | 0.243 | 0.15 |
|  |  | CCR2 | 399 | 14.013 | 0.115 (-0.108) | 0.142 | 0.10 |
|  |  | CXCR3 | 199 | 14.853 | 0.191 (-0.032) | 0.143 | 0.13 |
| CXCR3 | 994 | CXCR3 | 199 | 4.727 | 0.322 | 0.198 | 0.15 |
|  |  | CCR2 | 399 | 13.681 | 0.125 (-0.197) | 0.124 | 0.13 |
|  |  | CCR3 | 121 | 8.029 | 0.182 (-0.140) | 0.142 | 0.13 |

**Table 4.** Performance of Keras/TensorFlow NN in the cannabinoid receptor subtype selectivity tasks.

| Training set | Number of datapoints | Validation set | Number of datapoints | Loss | Accuracy (change) | Average Tanimoto coefficient<br>training vs. validation set |
| --- | --- | --- | --- | --- | --- | --- |
| 2023 ChEMBL datasets |  |  |  |  |  |  |
| CB1 | 4509 | CB1 | 902 | 2.455 | 0.503 | 0.129 |
|  |  | CB2 | 818 | 7.174 | 0.246 (-0.257) | 0.133 |
|  |  | CB2 selective | 35 | 6.499 | **0.0146** | 0.139 |
| CB2 | 4087 | CB2 | 818 | 2.987 | 0.418 | 0.137 |
|  |  | CB1 | 902 | 5.354 | 0.291 (-0.127) | 0.131 |
|  |  | CB2 selective | 35 | 2.450 | **0.521** | 0.141 |
| 2020 ChEMBL datasets [83] from https://db-gpcr.chem.uw.edu.pl |  |  |  |  |  |  |
| CB1 | 1566 | CB1 | 314 | 1.943 | 0.464 | 0.152 |
|  |  | CB2 | 418 | 4.417 | 0.203 (-0.261) | 0.148 |
|  |  | CB2 selective | 35 | 9.187 | **0.0135** | 0.150 |
| CB2 | 2093 | CB2 | 418 | 1.919 | 0.509 | 0.152 |
|  |  | CB1 | 314 | 4.263 | 0.210 (-0.299) | 0.147 |
|  |  | CB2 selective | 35 | 0.487 | **0.876** | 0.153 |

These results confirmed that although the composition of the training dataset has a noticeable impact on the classification results [63], neural networks are still able to classify correctly despite increase of noise in the training sets. Here, noise in datasets was introduced by adding inactive compounds to the current dataset. This advantage of NNs over supervised methods like gradient boosting decision trees (LightGBM) is mostly due to the fact that NNs can also act as unsupervised learners using unlabeled datasets for training. What is more, adding inactive or weakly active compounds to training sets only slightly worsened the accuracy of the activity prediction, which was also expected based on [54]. Adding inactive compounds to training sets could improve the binary classification (active vs. inactive compounds) but not the activity value prediction, which is a multiclass classification task [54].

If we compare the results presented in Table 3 and 4, the NN model trained on the cannabinoid receptor datasets seems to be more accurate in the selectivity prediction than models trained on the chemokine receptor datasets. In the case of the CB1 model, the prediction accuracy dropped by more than 0.2 when the validation set with the inconsistent receptor subtype was tested. In the case of the CB2 model, the accuracy changed even more—by 0.3. In the case of the chemokine receptor models, the most significant change in the accuracy was for the CXCR3 model (nearly 0.2 for the CCR2 validation set), but the remaining models showed only ca. 0.1 or less change in the accuracy. The worst model regarding the selectivity prediction was the CCR2 model, which is consistent with the fact that the CCR2 ligands from the training set were almost as similar to ligands from the CCR2 validation as from the CCR3 or CXCR3 validation sets. The CXCR3 model performed the best in the receptor subtype selectivity prediction task also for the same reason. The CXCR3 ligands retrieved from ChEMBL were the most dissimilar to both CCR2 and CCR3 ligands. In all cases, the prediction accuracy of NNs correlated with values of the Tanimoto coefficient between the training and validation sets.

## 3. Discussion and Conclusions

Due to the role they play in numerous diseases, chemokine receptors represent promising drug targets—however, drug design is hindered by the unavailability of many of their structures. In such cases, homology modeling makes it possible to create models of receptors based on their similarity to other receptors with solved structures. Though this can be done using webservers, standalone programs, such as Modeller, give researchers the opportunity to take a more hands-on approach and adjust the modeling process to suit their own needs. The created models can then be used in SBVS in order to search for novel active compounds for the receptors in question, and the results validated through the use of properly trained machine learning algorithms. Regarding virtual screening, the comparison to known CXCR3 ligands showed that our recently developed machine learning approach to ligand-based virtual screening provides substantially new information on the compound activity, different to predictions made by molecular docking in SBVS. Nevertheless, Keras/TensorFlow NN or LightGBM cannot be used solely but rather as a filtering method to decrease the number of compounds tested in SBVS for their precise binding modes and affinities. Machine learning also allows to screen much larger compound libraries than those accessible to SBVS. Such novel algorithms offer better accuracy and better computational time efficiency than classical QSAR methods. The pre-filtering of large compound libraries before the SBVS step requires accurate but fast computational methods, which can be easily fulfilled by ML. In our opinion, the only limitation of ML remains in its dependency on the composition of the used training datasets [54]. This seems to be more crucial for LightGBM, while NNs encounter problems arising from the limited size of the assay-derived datasets.

Molecular dynamics, though more computationally expensive than LBVS or SBVS, is much more reliable than these methods in the validation of the ligand-receptor interactions, as it provides a dynamic image of the protein system in a time-dependent manner. Here, MD simulations allowed to decide which of the previously selected compounds could serve as novel scaffolds for each of the studied receptors, and which would require modifications to improve their binding affinity. As a result, we obtained four novel chemotypes for CCR2, five for CCR3, and five for CXCR3. These molecules can serve as a basis for further drug design involving ligand binding assays and bioassays to confirm their ability to enhance the biological response of the receptor.

The combination of various computational methods allows to overcome the limitations of each method. For example, SBVS does not use any prior knowledge about known active ligands of a given target and encounters problems arising from a simplification of used force fields. Nanosecond MD simulations do not allow for scanning of all possible receptor binding sites and all possible ligand conformations. Machine learning used in LBVS does not use any explicit information about the receptor and its interactions with ligands. On the other hand, SBVS allows to perform an exhaustive search through all possible ligand conformations and ligand-receptor interactions to find the global free energy minimum. Nanosecond MD simulations allow unstable ligand-receptor interactions to be discarded and ligand binding modes to be corrected using detailed all-atom force fields. ML can perform an extremely fast search for active ligands among huge datasets of compounds and thus significantly limits the number of ligands to be tested in SBVS. GPU-accelerated neural networks designed in Keras/TensorFlow or using GPUs for LighGBM offer the next level of processing cheminformatic data.

Among ML methods, NNs or deep learning NNs built on the Keras/TensorFlow platform have been used so far mainly in binary classification tasks in drug design [91]. Here, we showed that NNs can also be used in drug design as efficient multi-class classifiers when trained on the datasets with discrete compound activity values [54]. To our knowledge, this is the first such application of Keras/TensorFlow NNs. Keras/TensorFlow NN multiclass classifier allows to discard not only inactive compounds from active ones, but also low-active compounds from highly active compounds. This is especially important for drug design referring to large datasets, in which the number of low-active compounds is so high and they are so diverse that they would introduce nothing but noise when used as training sets for binary classification.

Another important application of NN models is the prediction of the receptor subtype selectivity of a compound. As we showed, Keras/TensorFlow NNs can accurately distinguish ligand datasets matching different receptor subtypes. The only requirement is a sufficient dissimilarity between such ligand datasets, which was met in the case of CB1/CB2 datasets. Structural differences between ligands of different chemokine receptor subtypes were hardly sufficient, except for the CXCR3 dataset. Thus, based on the datasets currently available in ChEMBL we could only develop the CB1/CB2 selective NN model and CCR/CXCR selective model of an accuracy sufficient for drug design purposes.

## 4. Materials and Methods

### 4.1. Ligand-based Virtual Screening

In the LBVS step, we used a method described in detail elsewhere [54]. The method uses the Keras/TensorFlow library for constructing, training, and evaluating the currently used sequential model of neural network. NN was trained on the ChEMBL datasets of CCR2, CCR3, and CXCR3 compounds following procedure described in [54]. Extended connectivity fingerprints with bond diameter 4 (ECFP4) [77] based on Morgan fingerprints were used to describe compound features with RDKit [76]. To emphasize, Keras/TensorFlow NN was used here not as a typical binary NN classifier but as a multiclass classifier that is able to distinguish not only active and inactive compounds, but also low and medium-active from highly active compounds. This was done by labeling the datasets with seven activity categories based on logarithmic pChEMBL values: 1 (below 4), 2 (4–5), 3 (5–6), 4 (6–7), 5 (7–8), 6 (8–9), and 7 (above 9). Categories 5, 6 and 7 referred to highly active compounds, while 3 and 4 to medium-active, and 1 and 2 to inactive or low-active compounds. An NN was built and trained using the categorical cross-entropy loss function, stochastic gradient descent to minimize the loss function (Adaptive Moment Estimation Optimizer). Due to the multi-class application of NN, Softmax conversion leading to a probability distribution was used as the activation function for the last layer, instead of the sigmoid function that is used typically for binary classification. The Rectified Linear Unit activation function (ReLU) was used for hidden layers for quick convergence. 1000 epochs were used to ensure the sufficient minimization of the model, although a much smaller number could be also used, e.g., 200, as cross-entropy loss and accuracy stabilized after 200 epochs (see Appendix S1, Figure S2-S3).

In principle, in the case of neural networks fitted to solve big data problems, increasing the training set from 40 % to 80 % (see Appendix S1 Figure S4-S6) should improve both the model accuracy and the model training efficiency. This improvement was indeed visible in the case of CCR2 and CCR3 (Appendix Figure S4). Nevertheless, the bootstrappping analysis should be performed to undoutedly confirm this.

For the receptor subtype selectivity tests, the following curated datasets were used for training (80 % randomly selected compounds from the ChEMBL-retrieved datasets): 1995 (CCR2), 603 (CCR3), 994 (CXCR3), 4509 (CB1), 4087 (CB2), and for validation the remaining compounds were used. For cannabinoid receptors, two additional training sets [63,83] from https://db-gpcr.chem.uw.edu.pl were used, consisting of 1566 and 2093 compounds (CB1 and CB2, respectively). A further 35 CB2-selective compounds (from https://db-gpcr.chem.uw.edu.pl) were used as one of the validation sets included in Table 4. To generate the results presented in Table 4, the number of epochs were set to 100, and the average loss and accuracy was computed for 100 independent training runs of NN.

Python scripts with imported modules from the latest versions of RDKit, scikit-learn, Keras, and Tensorflow were used for data processing.

### 4.2. Preparation of CCR2, CCR3, and CXCR3 structures

The 6GPX structure [92,93] of the inactive-state CCR2 receptor was downloaded from the Protein Data Bank (PDB) [94], and a model of CCR3 was generated using the Robetta webserver [95]. Both the structure and the model were preprocessed using Maestro [96] and evaluated as described in a previous study [54].

The amino acid sequence of CXCR3-A was obtained from UniProt [97] (see Appendix S1 Fig. S1). Protein BLAST was used to perform a search against PDB to find solved GPCR structures with a high sequence similarity to CXCR3-A [98]. Of these, the inactive-state 5LWE [99,100] and 6MEO [101,102] PDB entries, with a 33.44% and 36.49% sequence similarity to the target, respectively, were selected as templates for homology modeling. Modeller [103] was used to generate 5000 models of CXCR3-A. The lowest energy models were analyzed in PyMol [104] and validated by comparing their structures to those of other GPCRs with well-described molecular switches, including PDB entries 7O7F [105,106] and 6WWZ [107,108].

### 4.3. Structure-based Virtual Screening

SBVS assisted by machine learning was performed with Glide for CCR2 and CCR3, and described previously [54]. Here, to confirm their binding modes and to test if two molecular docking programs based on completely different force fields (OPLS and Amber for Glide and AutoDock, respectively) provide similar results we used AutoDock Vina [43,44,109]. Although the compound ranking proposed by AutoDock Vina was very similar to the one obtained previously by Glide, a few compounds were discarded due to significant differences in their binding modes provided with AutoDock Vina in comparison to Glide results. The remaining CCR2 and CCR3 compounds were subjected to validation with MD simulations.

The validated model of CXCR3 was used for structure-based virtual screening (SBVS) with AutoDock Vina, using the Enamine Hit Locator Library (HLL) [110], consisting of over 460 000 compounds. The position of the grid box for AutoDock Vina was determined based on the positions of the ligands in the corresponding template structures, and its size was 31.19×29.17×38.56. Ten binding modes were generated for each ligand, and the energy cut-off for selecting ligand poses was equal to -10.5. The results were analyzed using the *vs-analysis.py* script [111], and 31 compounds with the best binding affinities were selected for further investigation.

A set of known CXCR3 inhibitors—the IC50 subset—was downloaded from the ChEMBL (accessed: May 2023). After the data was curated and compounds with no specified activity values (pChEMBL values) were removed, the CXCR3 dataset was used as a training set for a neural network implemented in Keras/TensorFlow according to a procedure described elsewhere [54]. The algorithm was then used to predict the activity values of the molecules in the HLL compound library. The compounds with the highest predicted activity values (above 9) were mapped against those obtained via SBVS, and as a result, nine potential CXCR3 actives were obtained. Out of these, five the best-assessed compounds were selected for further MD simulations.

### 4.4. Molecular Dynamics Simulations

For the selected compounds, their complexes with receptors for the MD simulations were prepared using CHARMM-GUI’s [112–114] Membrane Builder [115–118]. Information about the disulphide bonds in the receptor structures was provided based on known structures of chemokine receptors in the PDB and the ligand parameterization was performed using CGenFF [119] and 3D structural files generated by Maestro. The ligand-receptor complexes were inserted into a lipid bilayer consisting of a 3:1 ratio of POPC (1-palmitoyl-2-oleoyl-sn-glycero-3-phosphocholine) to cholesterol. The periodic rectangular water box (TIP3P) was fitted to the complex and each simulation system was neutralized by adding Na^+^ and Cl^-^ ions at a concentration of 0.15 M. The number of atoms in each simulation system was equal to between 135000 and 148000 atoms, depending on the system. The Charmm36 force field was used in each simulation.

The equilibration step included 10,000 steps of the steepest descent minimization, then 25,000 steps of the conjugated gradients minimization. The equilibration simulation was performed in NVT using the Langevin dynamics (303.15 K). The time integration step in the equilibration and production runs was set to 2 fs. The production run in NPT was performed using the Langevin piston Nose-Hoover method (1 bar, 303.15 K) and lasted for 100 ns for each system. The GPU-accelerated version of NAMD [120] was used for all MD simulations. The obtained trajectories were analyzed using VMD [121].

### 4.5. Suggested Structural Modification of Active Compounds

Chemical modifications of functional groups of the proposed active compounds for each receptor were suggested in order to improve their binding affinities. Maestro was used to analyze the interactions between the modified ligands and the receptor in the final frame of the MD simulation and to suggest possible changes. Modified structures of proposed compounds were minimized in Maestro (OPLS4 force field), in order to prevent clashes.

### 4.6 Structural Comparison of CXCR3 Antagonists

Compounds described by Meyer et al. [32] were reproduced in Maestro in order to perform a search for similar structures in the HLL compound library. The Fingerprint Similarity tool was used with the Tanimoto similarity metric. The docking scores and predicted activities as well as their ranks provided by Keras/TensorFlow NN were extracted for compounds with the highest Tanimoto coefficients.

## Supplementary Materials

The following supporting information can be downloaded at: www.mdpi.com/xxx/s1

## Author Contributions

Conceptualization, P.D., D.L.; Methodology, P.D., K.J., D.L.; Software, P.D., K.J., D.L.; Validation, P.D., K.J., D.L.; Formal Analysis, P.D., K.J., D.L.; Investigation, P.D., K.J., A.A., D.L.; Resources, D.L.; Data Curation, P.D., D.L.; Writing—Original Draft Preparation, P.D., K.J., A.A., D.L.; Writing—Review & Editing, P.D., A.A., D.L.; Visualization, P.D., D.L.; Supervision, D.L.; Project Administration, D.L.; Funding Acquisition, D.L.

Detailed contributions: Introduction - P.D., D.L., with contributions from A.A. (Table 1) and K.J.; Results - P.D. (Fig. 1–4, Table 2, sections 2.1, 2.2, 2.3, 2.4), D.L. (Tables 2–4, sections 2.2, 2.4, 2.5); Discussion and Conclusions - P.D., D.L.; Materials and Methods - P.D. (sections 4.1, 4.2, 4.3, 4.4, 4.5, 4.6), D.L. (section 4.1) with contributions from K.J. (performing AutoDock Vina computations for CCR2, CCR3, and CXCR3, and ML computations for CXCR3); Appendix S1 - P.D. (Fig. S1, Tables S1–S6), D.L. (Fig. S2–S6, Tables S7–S10); Graphical Abstract: D.L., P.D..

**Institutional Review Board Statement:** Not applicable.

**Informed Consent Statement:** Not applicable.

**Data Availability Statement:** Datasets used for NN training are publicly available on: https://www.ebi.ac.uk/chembl/ and https://db-gpcr.chem.uw.edu.pl.

**Conflicts of Interest:** The authors declare no conflict of interest.

## Abbreviations

GPCR: G protein-coupled receptor
CCL: C-C motif chemokine ligand
CXCL: C-X-C motif chemokine ligand
CCL: C-C motif chemokine ligand
CCR: C-C motif conventional chemokine receptor
CXCR: C-X-C motif chemokine receptor
ACKR: atypical chemokine receptor
MSMP: prostate-associated microseminoprotein
SBVS: structure-based virtual screening
LBVS: ligand-based virtual screening
MD: Molecular Dynamics
HLL: Hit Locator Library
NN: neural network
ML: machine learning
Cryo-EM: cryogenic electron microscopy

**Disclaimer/Publisher’s Note:** The statements, opinions and data contained in all publications are solely those of the individual author(s) and contributor(s) and not of MDPI and/or the editor(s). MDPI and/or the editor(s) disclaim responsibility for any injury to people or property resulting from any ideas, methods, instructions or products referred to in the content.

