## Appendix 1 for "Keras/TensorFlow in Drug Design for Immunity Disorders"

### Appendix S1

#### Keras/TensorFlow in Drug Design for Immunity Disorders


Paulina Dragan[1], Kavita Joshi[1], Alessandro Atzei[2], Dorota Latek[1, *]

[1]Faculty of Chemistry, University of Warsaw, Pasteura 1, 02-903 Warsaw, Poland

[2]Department of Life and Environmental Science, Food Toxicology Unit, University of Cagliari, University Campus of Monserrato, SS 554, 09042 Cagliari, Italy.


**Figure S1.** Multiple sequence alignment of chemokine receptors CCR2, CCR3, and CXCR3. The most conserved residues are highlighted in yellow and used for residue numbering in the Ballesteros-Weinstein notation [1]. The alignment was performed using Clustal Omega [2].

```
CLUSTAL O(1.2.4) multiple sequence alignment


sp|P49682|CXCR3_HUMAN      MVLEVSDHQVLNDAEVAALLENFSSSYDYGENESDSCCTSPPCPQDFSLNFDRAFLPALY     60
sp|P41597|CCR2_HUMAN       -MLSTSRSRFIRNTNESGEEVTTFFDYDY----------GAPCHKFDVKQIGAQLLPPLY     49
sp|P51677|CCR3_HUMAN       ---------MTTSLDTVETFGTTSYYDDV----------GLLCEKADTRALMAQFVPPLY     41
                                    .  . :      .     *           .  * :     :   ::* **
                                    1.50                        2.50
sp|P49682|CXCR3_HUMAN      SLLFLLGLLGNGAVAAVLLSRRTALSSTDTFLLHLAVADTLLVLTLPLWAVDA-AVQWVF     119
sp|P41597|CCR2_HUMAN       SLVFIFGFVGNMLVVLILINCKKLKCLTDIYLLNLAISDLLFLITLPLWAHSA-ANEWVF     108
sp|P51677|CCR3_HUMAN       SLVFTVGLLGNVVVVMILIKYRRLRIMTNIYLLNLAISDLLFLVTLPFWIHYVRGHNWVF     101
                           **:* .*::**  *. :*:. :     *: :**:**::* *:::***:*   . . :***
                                                       3.50                        4.50
sp|P49682|CXCR3_HUMAN      GSGLCKVAGALFNINFYAGALLLACISFDRYLNIVHATQLYRRGPPARVTLTCLAVWGLC     179
sp|P41597|CCR2_HUMAN       GNAMCKLFTGLYHIGYFGGIFFIILLTIDRYLAIVHAVFALKARTVTFGVVTSVITWLVA     168
sp|P51677|CCR3_HUMAN       GHGMCKLLSGFYHTGLYSEIFFIILLTIDRYLAIVHAVFALRARTVTFGVITSIVTWGLA     161
                           * .:**:  .::: . :.  :::  :::**** ****.   :    :  .:*.: .* :.
                                                                             5.50
sp|P49682|CXCR3_HUMAN      LLFALPDFIFLSAHHDERLNATHCQYNFPQVGRT----ALRVLQLVAGFLLPLLVMAYCY     235
sp|P41597|CCR2_HUMAN       VFASVPGIIFTKCQKED--SVYVCGPYFPR----GWNNFHTIMRNILGLVLPLLIMVICY     222
sp|P51677|CCR3_HUMAN       VLAALPEFIFYETEELF--EETLCSALYPEDTVYSWRHFHTLRMTIFCLVLPLLVMAICY     219
                           :: ::* :** . ..    .   *   :*.           :   :  ::****:*. **
                                                             6.50
sp|P49682|CXCR3_HUMAN      AHILAVLLVSRGQR-RLRAMRLVVVVVVAFALCWTPYHLVVLVDILMDLGALARNCGRES     294
sp|P41597|CCR2_HUMAN       SGILKTLLRCRNEKKRHRAVRVIFTIMIVYFLFWTPYNIVILLNTFQEFF-GLSNCESTS     281
sp|P51677|CCR3_HUMAN       TGIIKTLLRCPSKK-KYKAIRLIFVIMAVFFIFWTPYNVAILLSSYQSIL-FGNDCERSK     277
                           : *: .** . .:: : :*:*::..:: .: : ****::.:*:.   .:     :*   .
                                              7.50
sp|P49682|CXCR3_HUMAN      RVDVAKSVTSGLGYMHCCLNPLLYAFVGVKFRERMWMLLLR-----LGCPNQRGLQRQPS     349
sp|P41597|CCR2_HUMAN       QLDQATQVTETLGMTHCCINPIIYAFVGEKFRSLFHIALGCRIAP-LQKPVCGGPGVRPG     340
sp|P51677|CCR3_HUMAN       HLDLVMLVTEVIAYSHCCMNPVIYAFVGERFRKYLRHFFHRHLLMHLGRYIP----FLPS     333
                           ::* .  **. :.  ***:**::***** :**. :   :       *          *.

sp|P49682|CXCR3_HUMAN      SSRR----------DSSW--SETSEASYS---GL368
sp|P41597|CCR2_HUMAN       KNVKVTTQGLLDGRGKGKSIGRAPEASLQDKEGA374
sp|P51677|CCR3_HUMAN       EKLE--------RTSSVSPSTAEPELSIVF----355
                           .. .          ..        * *
```

**Figure S2.** Training efficiency of Keras/TensorFlow NN using ChEMBL ligand datasets for chemokine receptors – values of the loss function. Here, 80% of the ChEMBL-retrieved datasets were used as training sets.

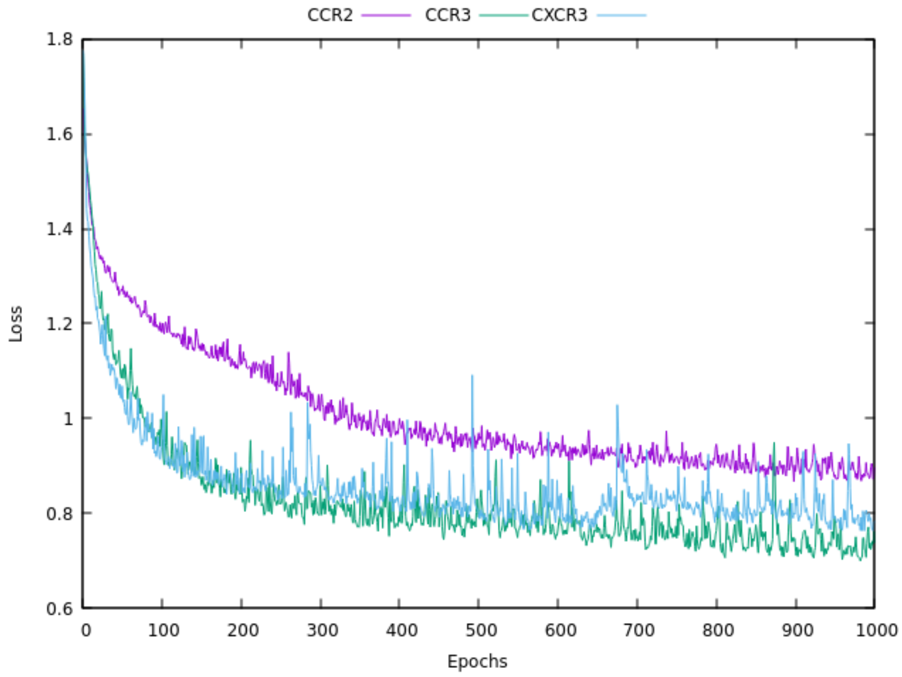


**Figure S3.** Training efficiency of Keras/TensorFlow NN using ChEMBL ligand datasets for chemokine receptors – the model accuracy. Here, 80% of the ChEMBL datasets were used as training sets.

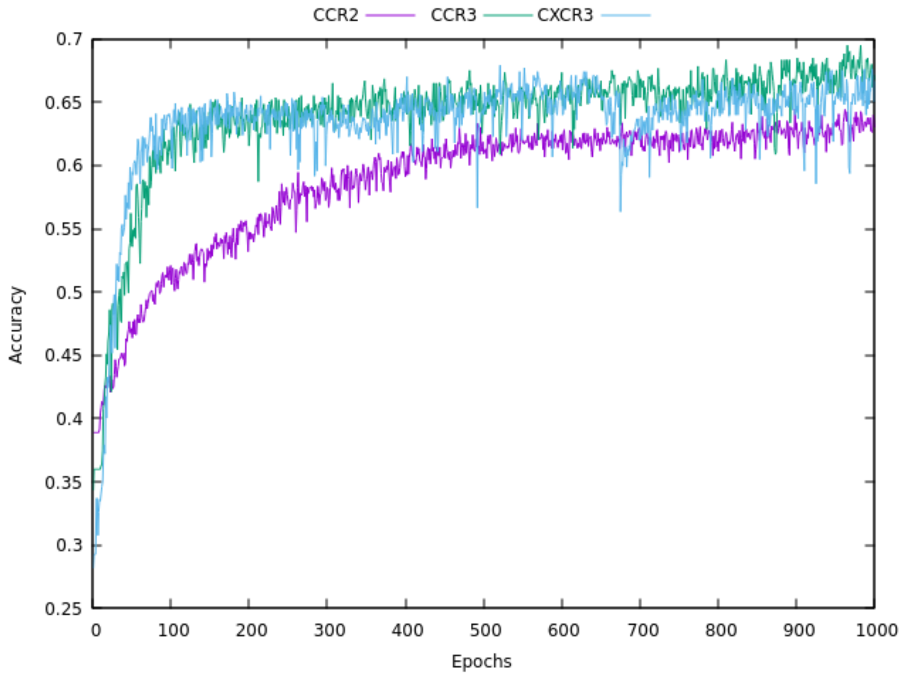

**Figure S4.** The impact of the quantity of the datasets on the model accuracy. Here, either 80% or 40% of the ChEMBL datasets were used as training sets.

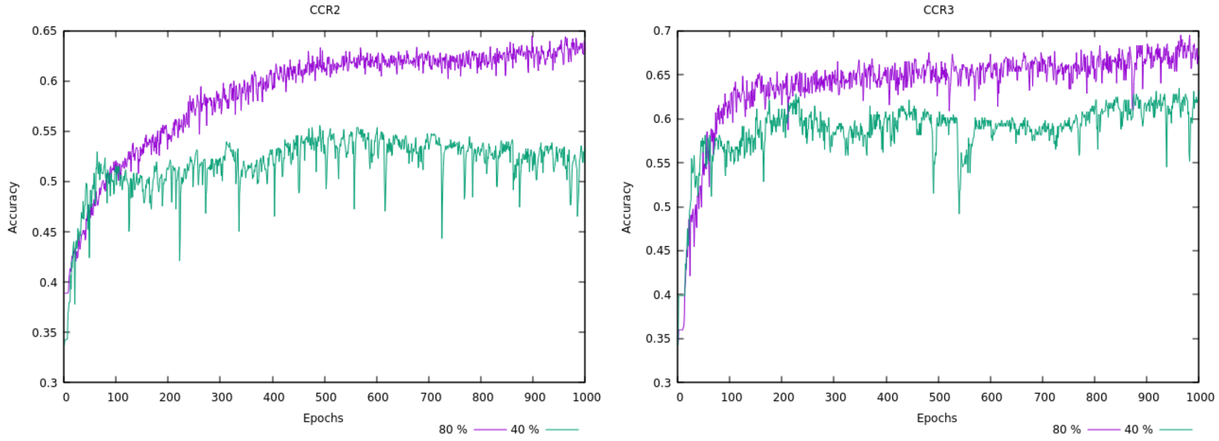


**Figure S5.** Training efficiency of Keras/TensorFlow NN using ChEMBL ligand datasets for the CB1 cannabinoid receptor. Here, 80% of the ChEMBL-retrieved datasets were used as training sets.

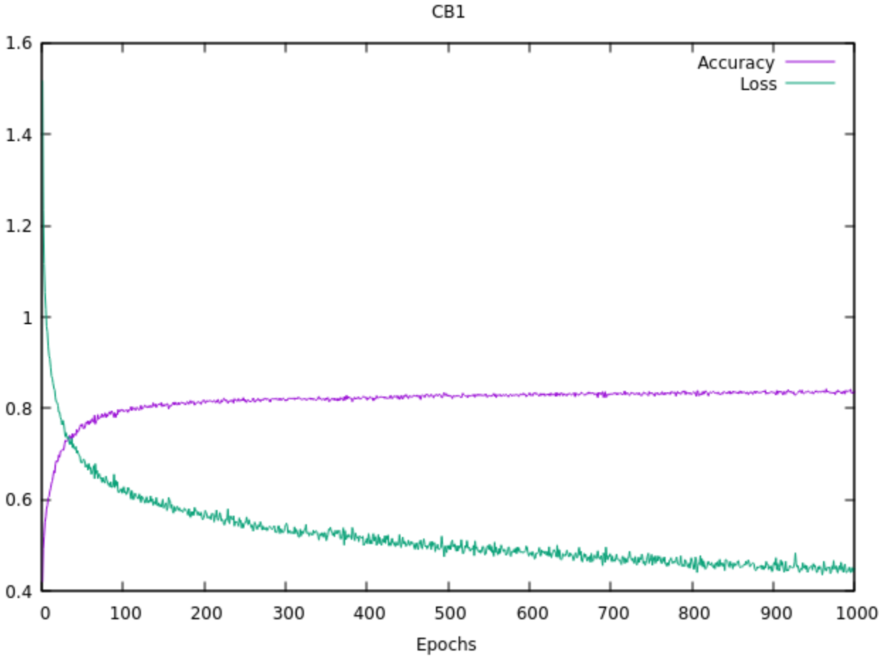

**Figure S6.** Training efficiency of Keras/TensorFlow NN using ChEMBL ligand datasets for the CB2 cannabinoid receptor. Here, 80% of the ChEMBL-retrieved datasets were used as training sets.

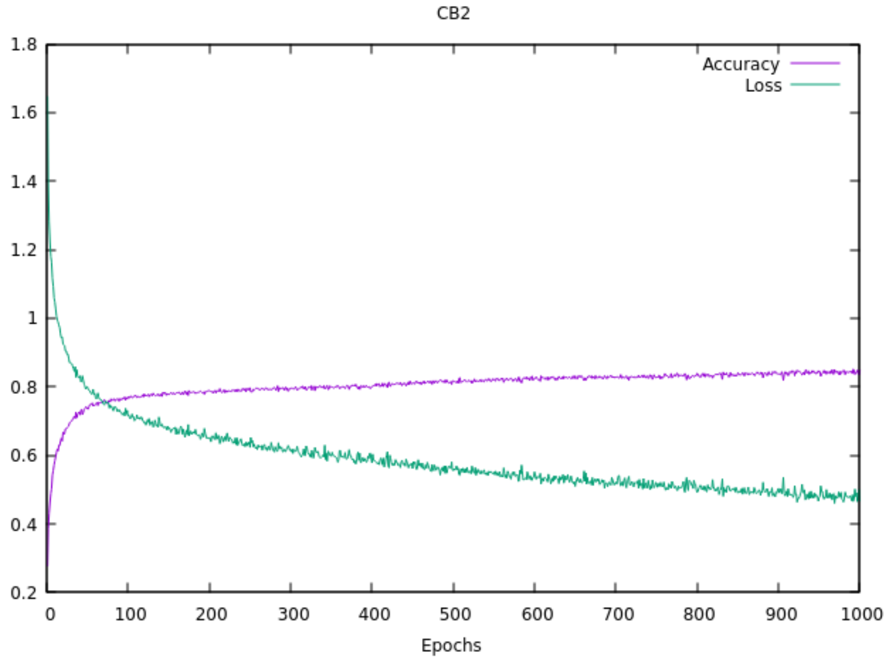

**Table S1**. The ligand-receptor interactions determined by Maestro [3] for the SBVS and MD results for CCR2.

| Compound id in Enamine HLL | SBVS-based ligand-receptor interactions | MD-based ligand-receptor interactions |
|---|---|---|
| Z144527132 | 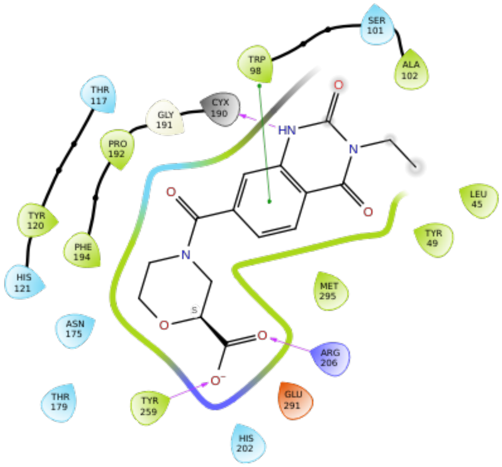 | 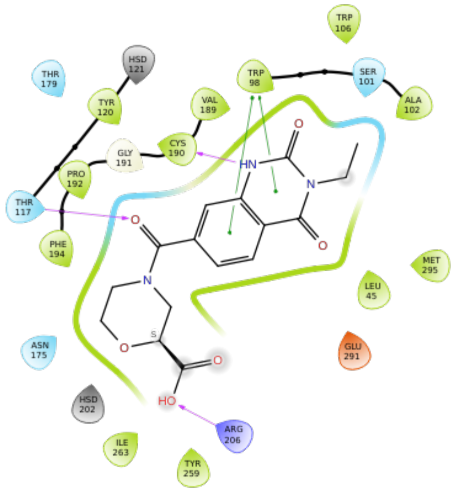 |
| Z199951150 | 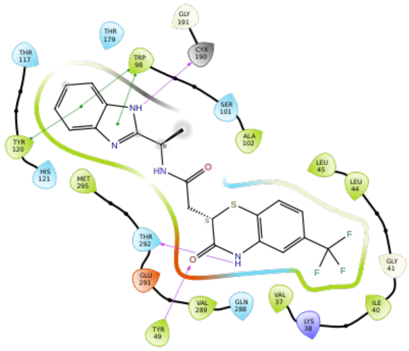 | 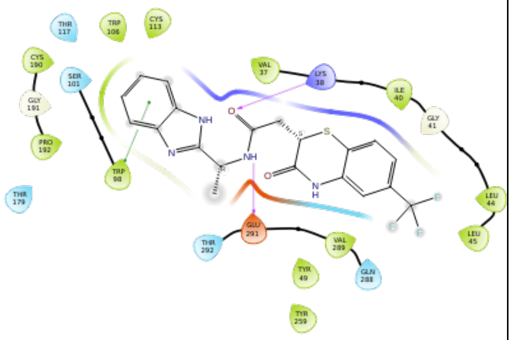 |

| | | |
|---|---|---|
| Z2607653068 | 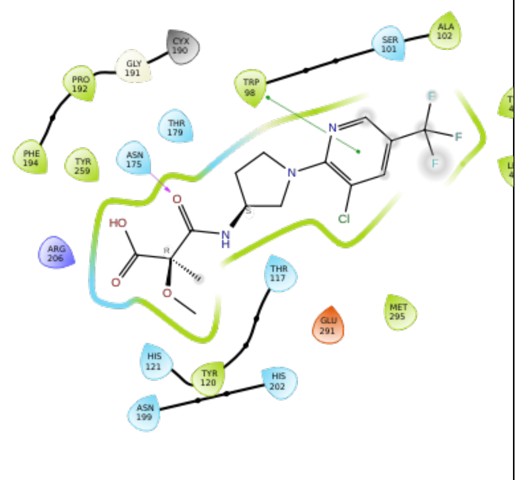 | 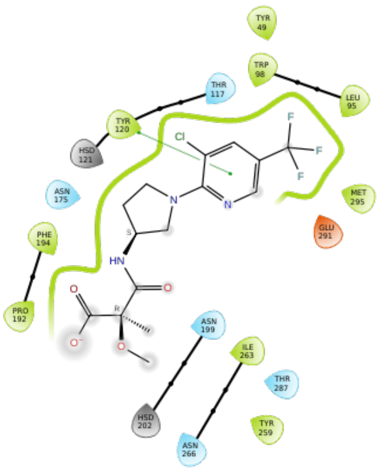 |
| Z45637008 | 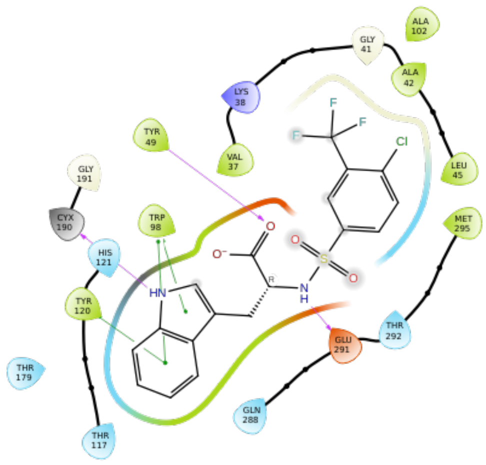 | 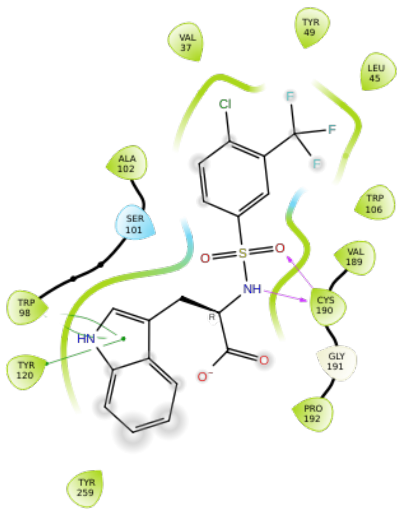 |

Charged (negative)
Charged (positive)
Glycine
Hydrophobic
Metal

Polar
Unspecified residue
Water
Hydration site
Hydration site (displaced)

Distance
H-bond
Halogen bond
Metal coordination
Pi-Pi stacking

Pi-cation
Salt bridge
Solvent exposure

**Table S2**. The ligand-receptor interactions determined by Maestro for the SBVS and MD results for CCR3.

| Compound id in Enamine HLL | SBVS-based ligand-receptor interactions | MD-based ligand-receptor interactions |
|---|---|---|
| Z1274732994 | 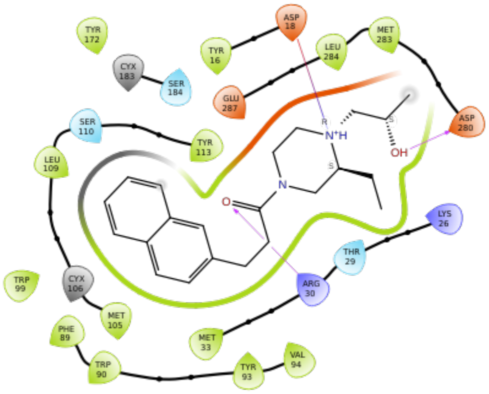 | 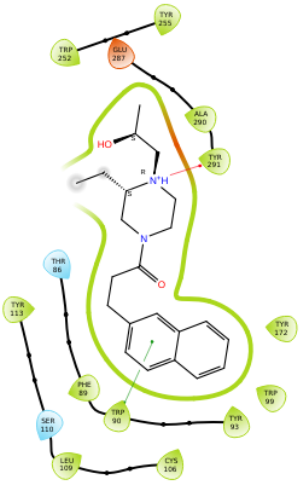 |
| Z1912507172 | 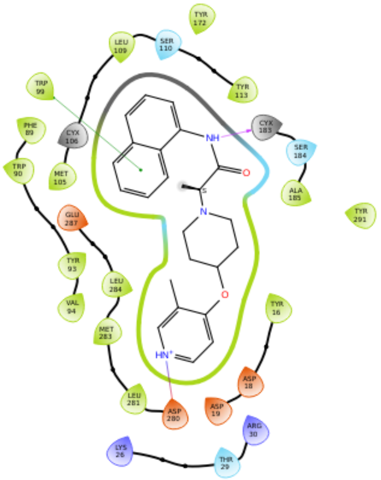 | 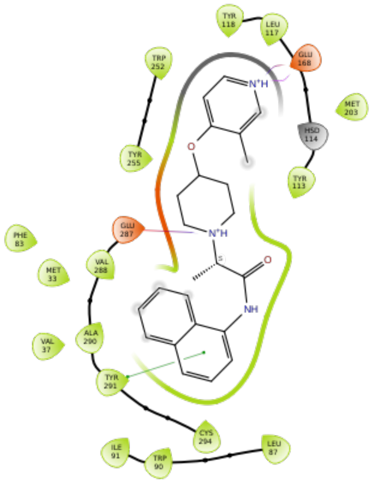 |

| | | |
|---|---|---|
| Z2441027668 | 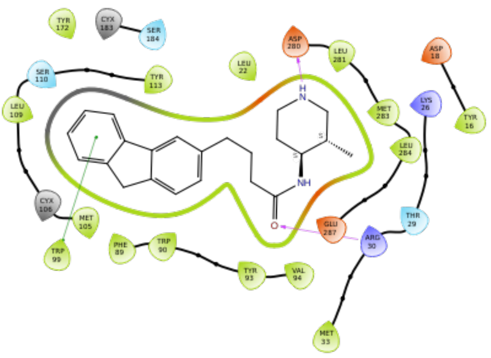 | 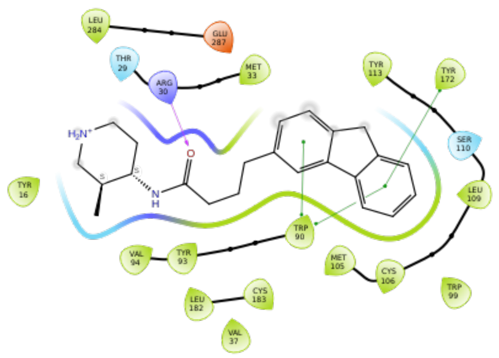 |
| Z2606182917 | 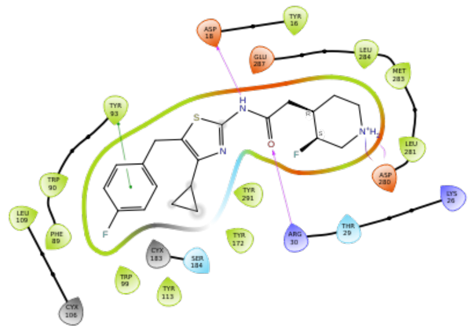 | 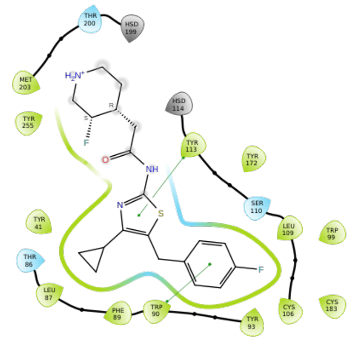 |
| Z2764968046 | 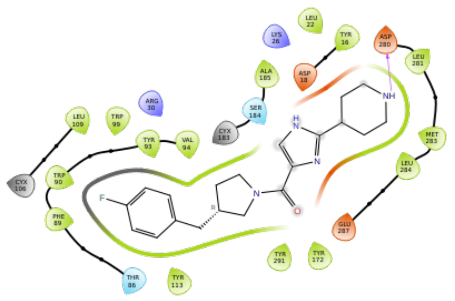 | 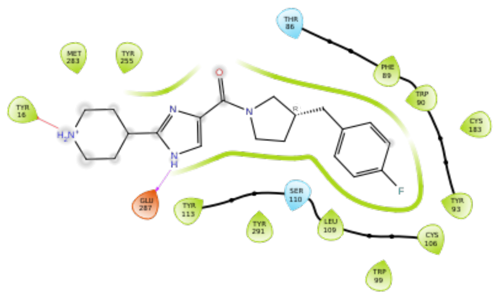 |

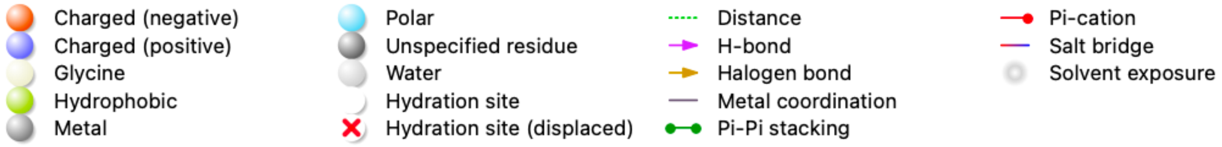

**Table S3.** The ligand-receptor interactions determined by Maestro for the SBVS and MD results for CXCR3.

| Compound id in Enamine HLL | SBVS-based ligand-receptor interactions | MD-based ligand-receptor interactions |
|---|---|---|
| Z107207944 | 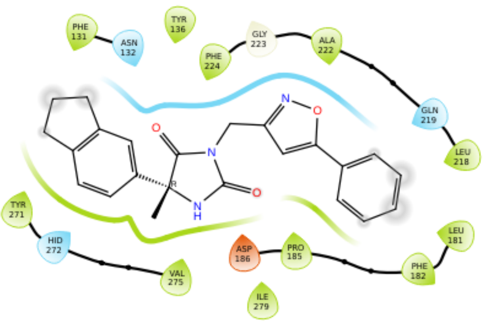 | 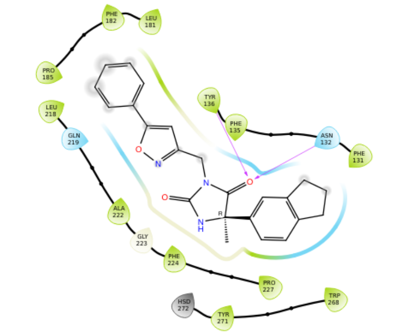 |
| Z1167188972 | 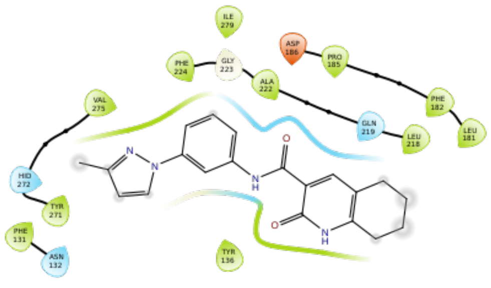 | 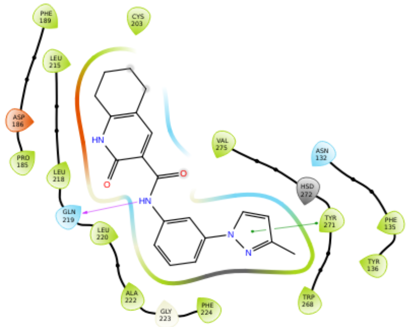 |
| Z1510954688 | 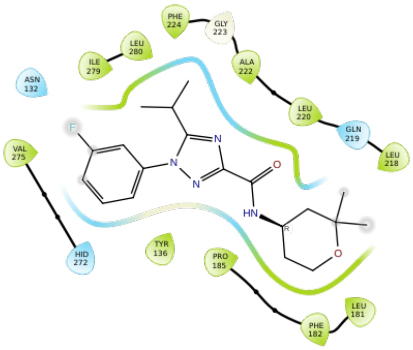 | 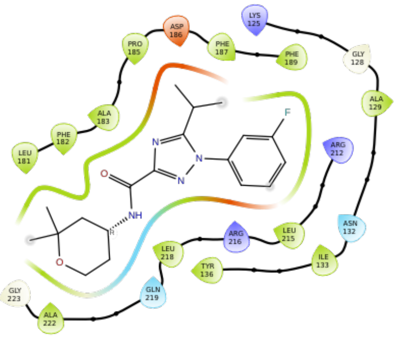 |

| | | |
|---|---|---|
| Z1903257002 | 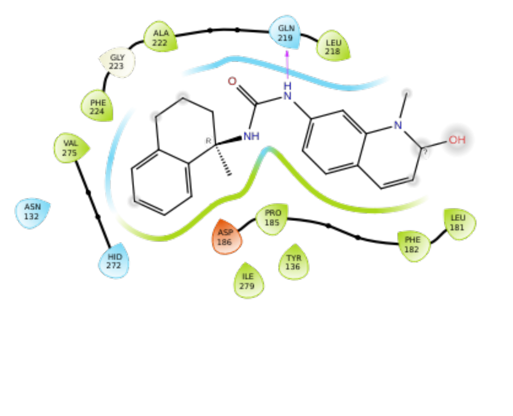 | 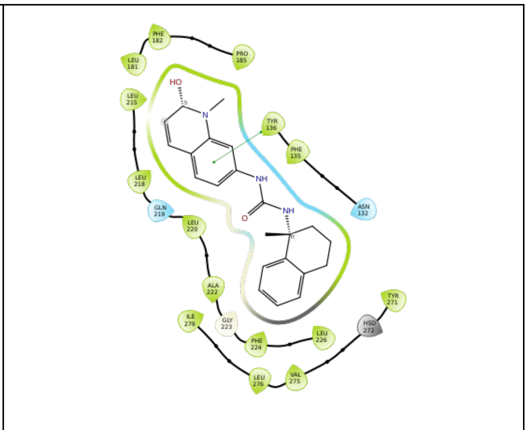 |
| Z2233592864 | 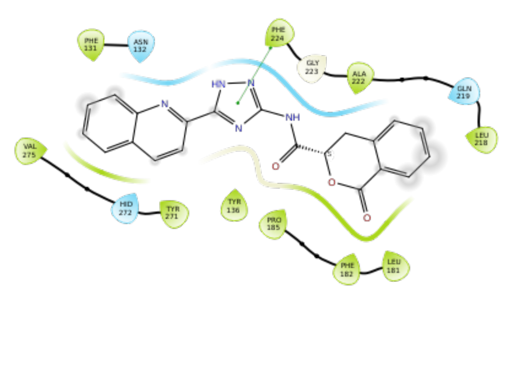 | 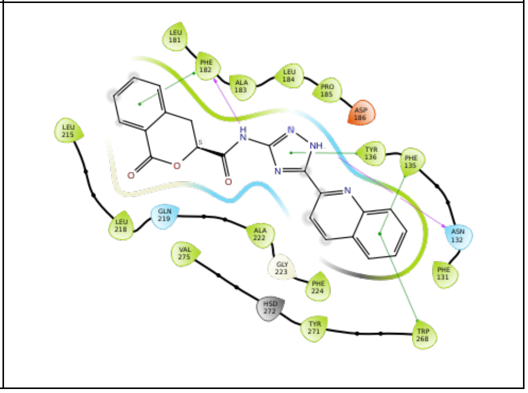 |

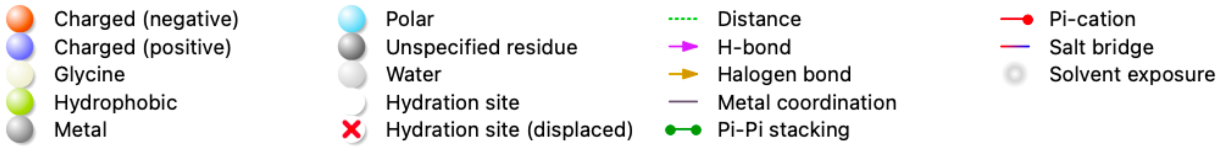

**Table S4.** Suggested modifications of the molecules selected for CCR2 and their contacts with the receptor predicted by Maestro. In the rightmost column: green dashed lines—good contacts, blue dashed lines—aromatic interactions, yellow dashed lines—hydrogen bonds.

| Original HLL compound | Modified structure of new compound | New interactions formed with the receptor | AutoDock Vina score |
|---|---|---|---|
| Z144527132 | 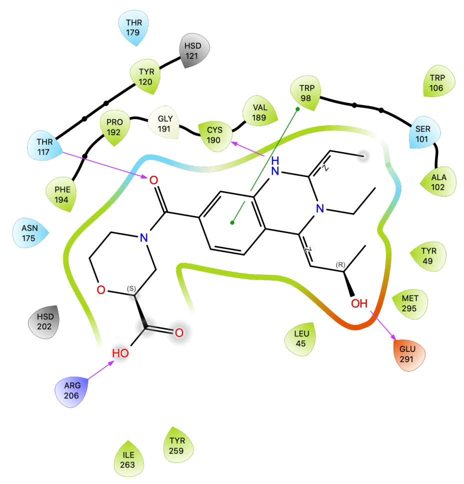  | 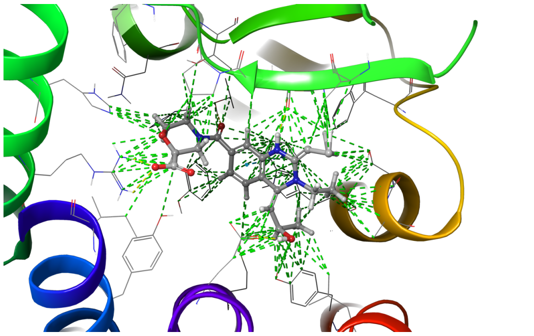 | -8.508 |
| Z199951150 | — | — | -9.305 |
| Z2607653068 | 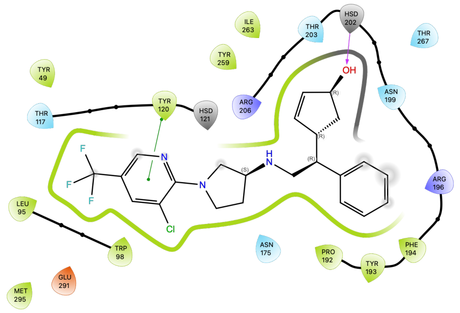  | 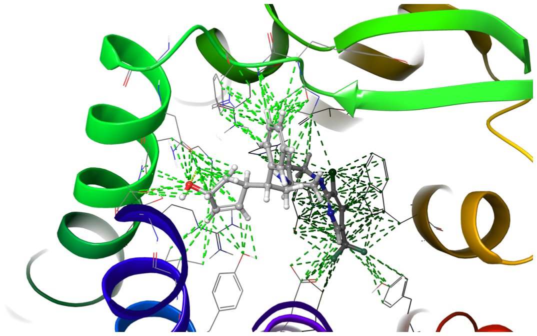 | -8.339 |
| Z45637008 | — | — | -8.103 |

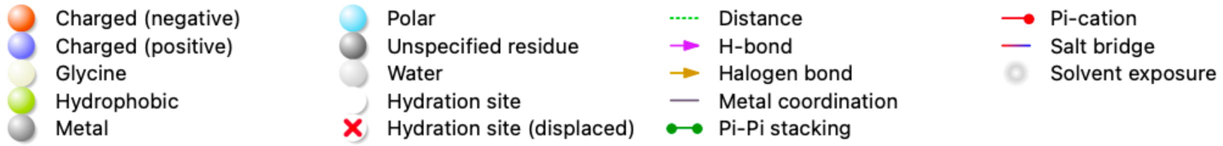

**Table S5**. Suggested modifications of the molecules selected for CCR3 and their contacts with the receptor predicted by Maestro. In the rightmost column: green dashed lines—good contacts, blue dashed lines—aromatic interactions, yellow dashed lines—hydrogen bonds, purple dashed lines—salt bridges, dark green—pi-cation interactions.

| Original HLL compound | Modified structure of new compound | New interactions formed with the receptor | AutoDock Vina score |
|---|---|---|---|
| Z1274732994 | 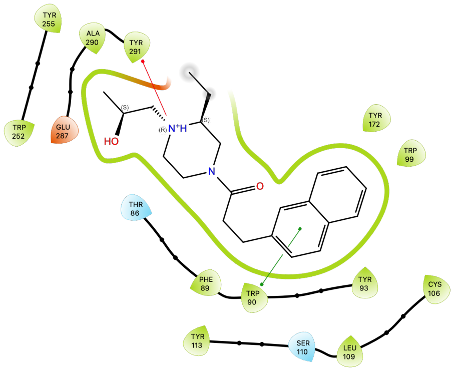  | 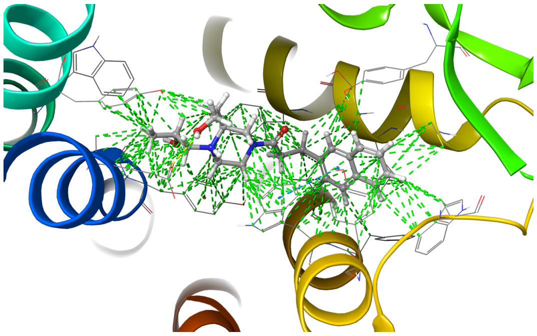 | -9.796 |
| Z1912507172 | 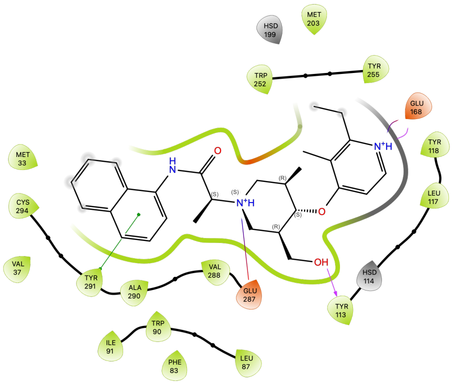  | 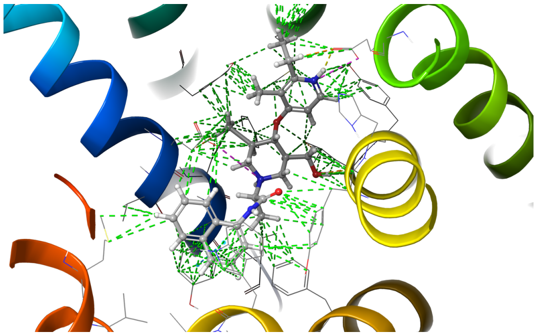 | -8.995 |

<table>
<tr><td>Z2441027668</td><td>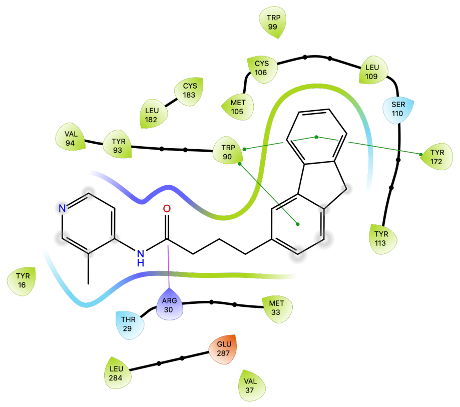
</td><td>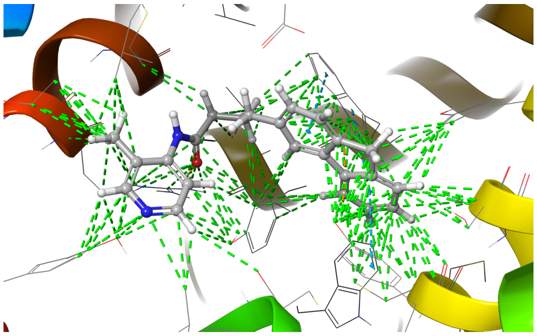</td><td>-7.486</td></tr>
<tr><td>Z2606182917</td><td>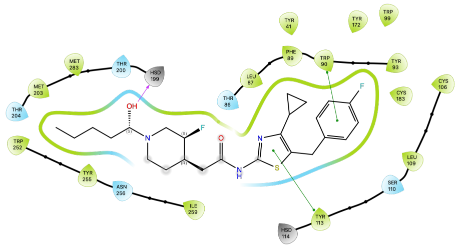</td><td>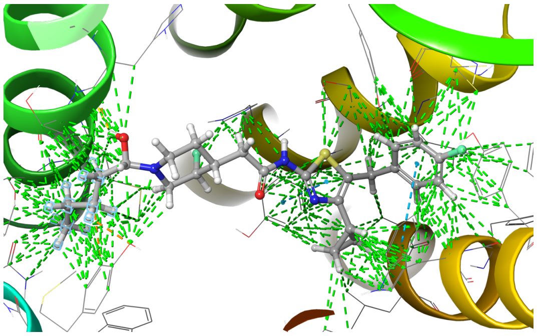</td><td>-8.387</td></tr>
<tr><td>Z2764968046</td><td>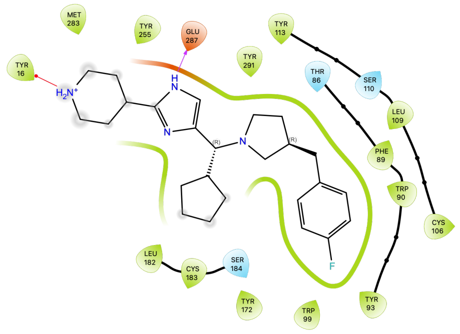
</td><td>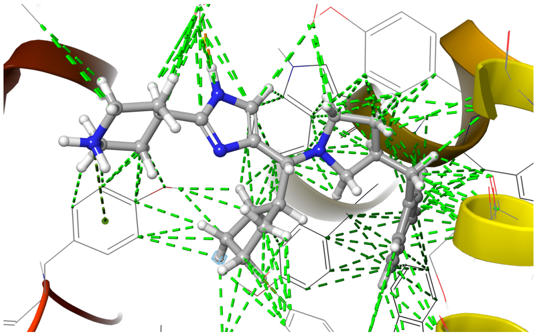</td><td>-8.926</td></tr>
</table>

Charged (negative)
Charged (positive)
Glycine
Hydrophobic
Metal
Polar
Unspecified residue
Water
Hydration site
Hydration site (displaced)
Distance
H-bond
Halogen bond
Metal coordination
Pi-Pi stacking
Pi-cation
Salt bridge
Solvent exposure

**Table S6**. Suggested modifications of the molecules selected for CXCR3 and their contacts with the receptor predicted by Maestro. In the rightmost column: green dashed lines—good contacts, blue dashed lines—aromatic interactions, yellow dashed lines—hydrogen bonds.

| Original HLL compound | Modified structure of new compound | New interactions formed with the receptor | AutoDock Vina score |
|---|---|---|---|
| Z107207944 | 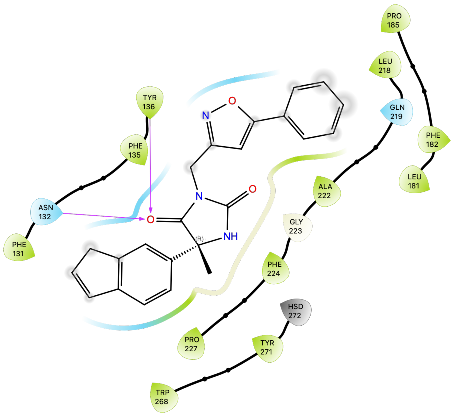 | 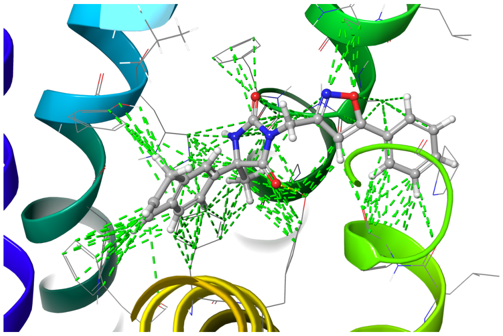 | -9.558 |
| Z1167188972 | 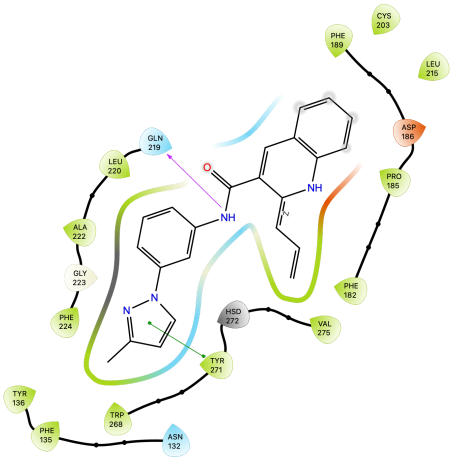 | 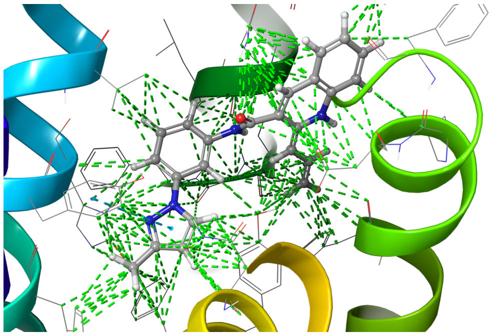 | -10.688 |

| | | | |
|---|---|---|---|
| Z1510954688 | 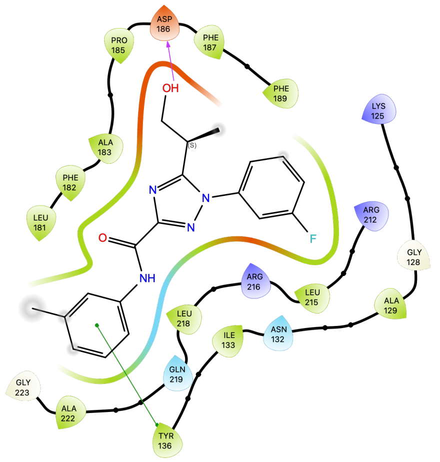  | 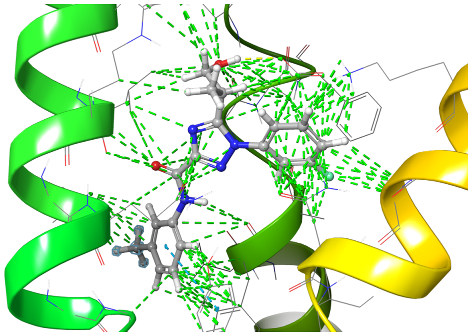 | -8.422 |
| Z1903257002 | 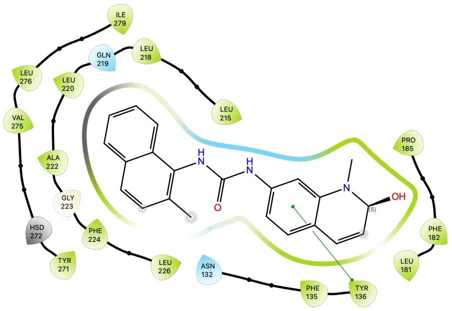  | 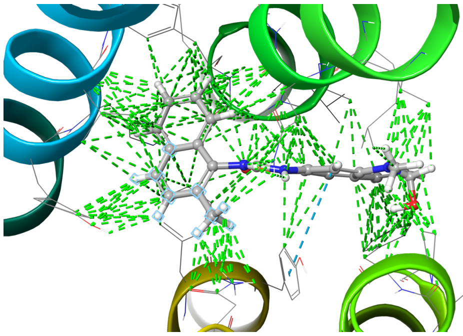 | -10.318 |
| Z2233592864 | — | — | -11.862 |

- Charged (negative)
- Charged (positive)
- Glycine
- Hydrophobic
- Metal
- Polar
- Unspecified residue
- Water
- Hydration site
- Hydration site (displaced)
- Distance
- H-bond
- Halogen bond
- Metal coordination
- Pi-Pi stacking
- Pi-cation
- Salt bridge
- Solvent exposure

**Table S7**. Binding modes obtained from AutoDock Vina for known CXCR3 antagonists. Polar contacts have been marked with yellow dashed lines with the respective receptor residues shown in sticks. Receptor structures were shown in the blue-to-red color scheme, while ligands were shown in blue.

| Known CXCR3 antagonist | AutoDock Vina prediction of its binding mode |
|---|---|
| ACT-672125 | 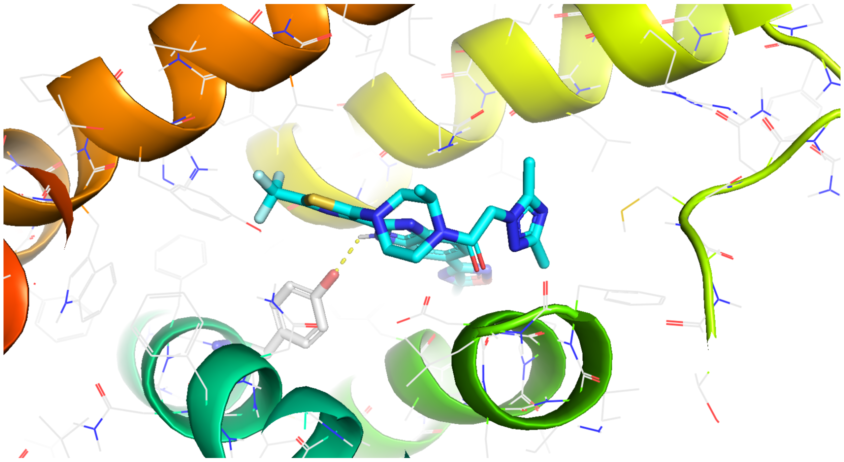 |
| ACT-660602 | 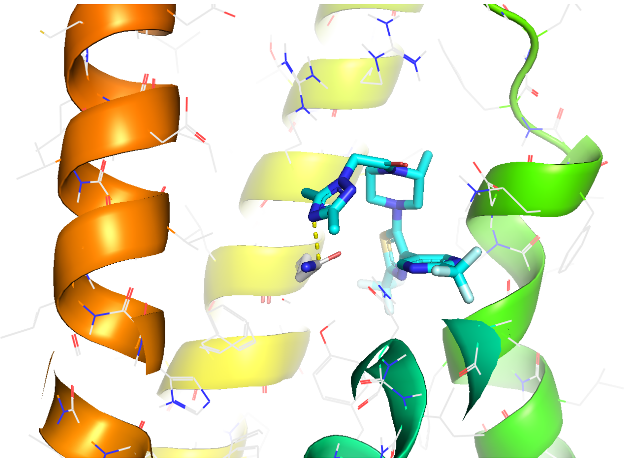 |

| | |
|---|---|
| ACT-777991 | 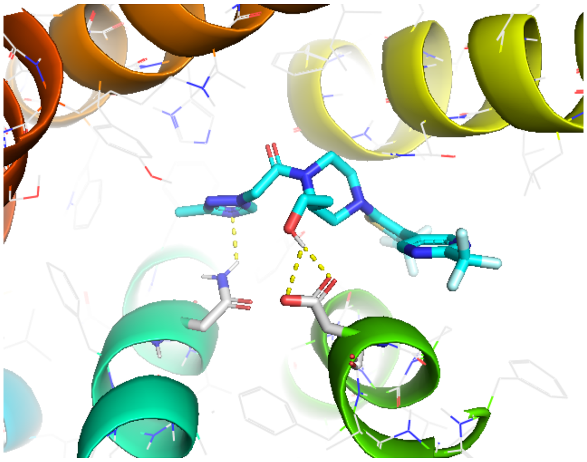 |
| VUF10661 | 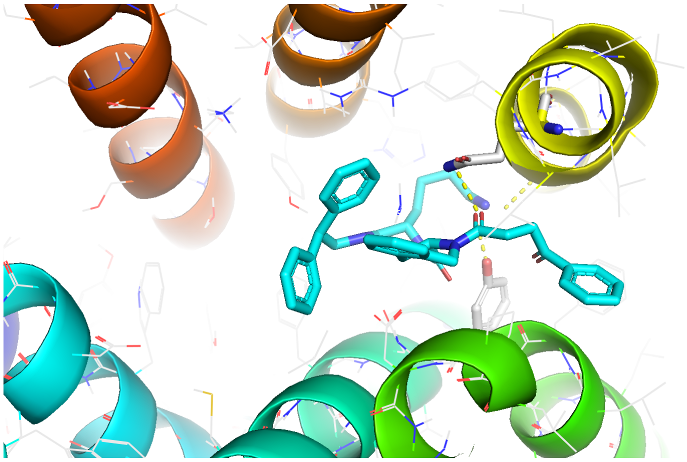 |

**Table S8.** Histograms showing the dataset classes distribution for the current and the previous CB1 and CB2 datasets (full datasets). PChEMBL values and corresponding classes used in training of NNs: 0–4 (class I), 4–5 (class II), 5–6 (class III), 6–7 (class IV), 7–8 (class V), 8–9 (class VI), above 9 (class VII).

<table>
<tr><td>Receptor</td><td>The current dataset</td><td>The previous dataset from https://db-gpcr.chem.uw.edu.pl</td></tr>
<tr><td>CB1</td><td>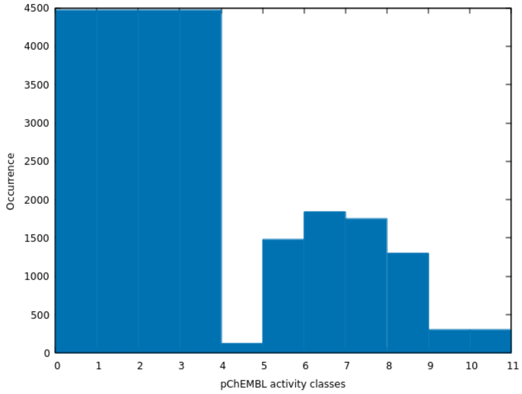
</td><td>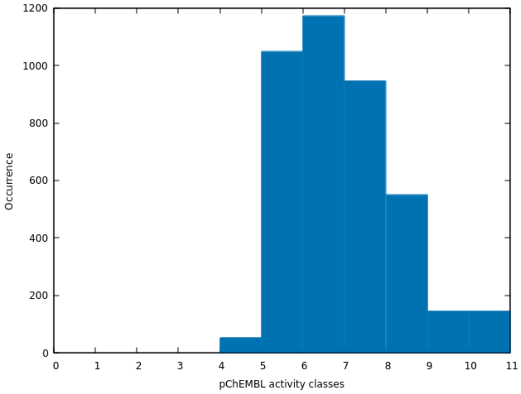
</td></tr>
<tr><td></td><td>Total of 5636 compounds</td><td>Total of 1958 compounds</td></tr>
<tr><td>CB2</td><td>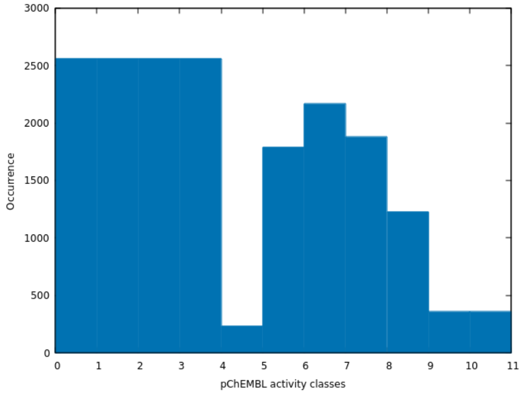
</td><td>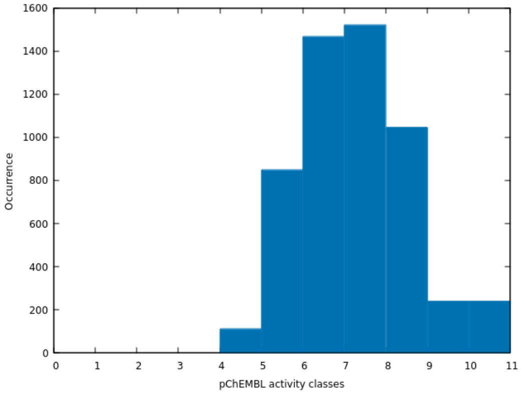
</td></tr>
<tr><td></td><td>Total of 5109 compounds</td><td>Total of 2616 compounds</td></tr>
</table>

**Table S9.** Histograms of Tanimoto coefficients between the training and validation datasets.

| Training set | Number of datapoints | Validation set | Number of datapoints | Histogram of Tanimoto coefficients<br><br>training vs. validation set |
|---|---|---|---|---|
| CCR2 | 1995 | CCR2 | **399** | 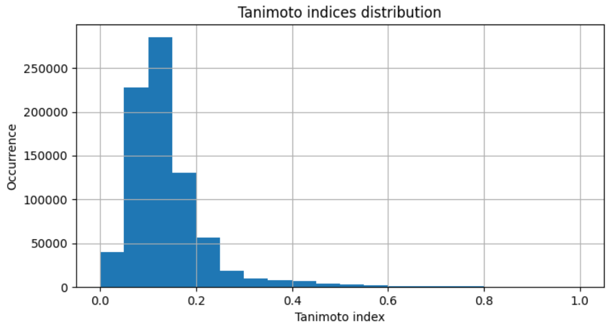  |
| | | CCR3 | 121 | 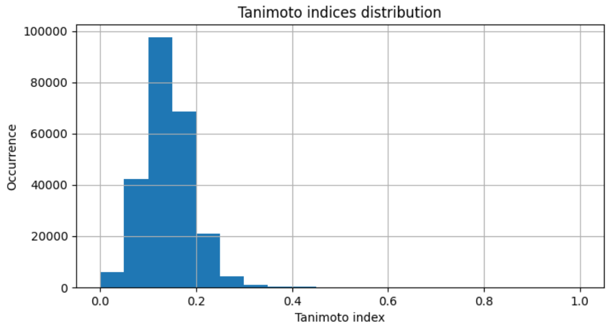  |
| | | CXCR3 | 199 | 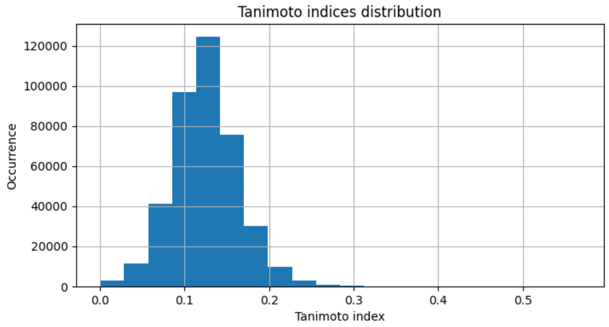  |
| CCR3 | 603 | CCR3 | 121 | 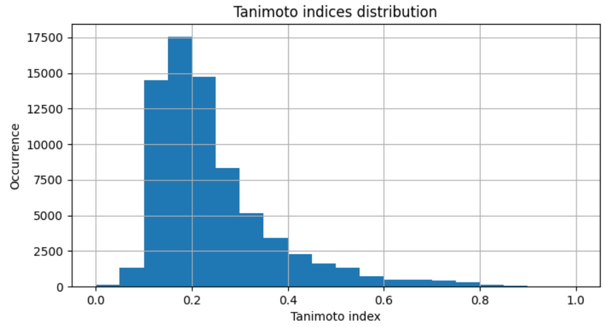  |

| | | | | |
|---|---|---|---|---|
| | | CCR2 | 399 | 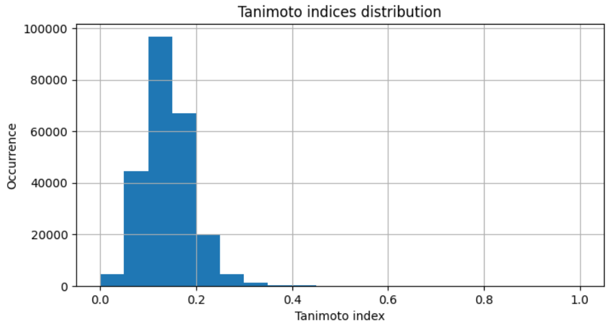 |
| | | CXCR3 | 199 | 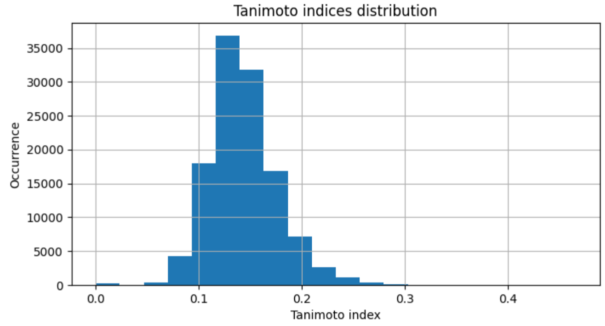 |
| CXCR3 | 994 | CXCR3 | 199 | 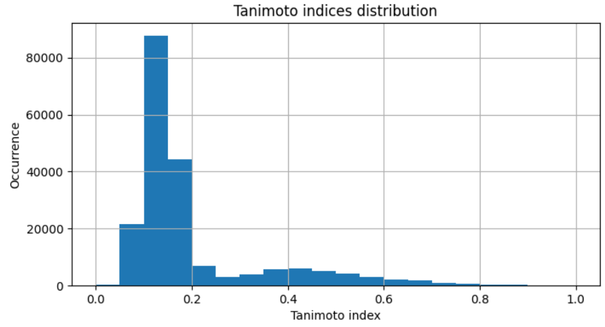 |
| | | CCR2 | 399 | 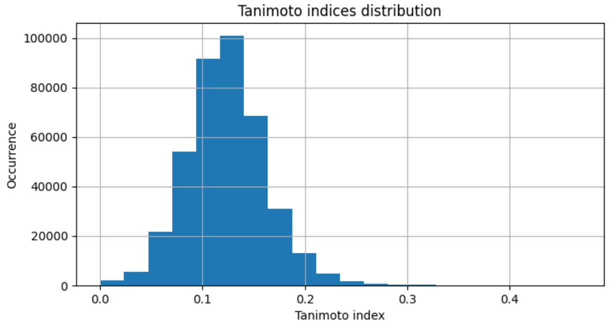 |

| | | | | |
|---|---|---|---|---|
| | | CCR3 | 121 | 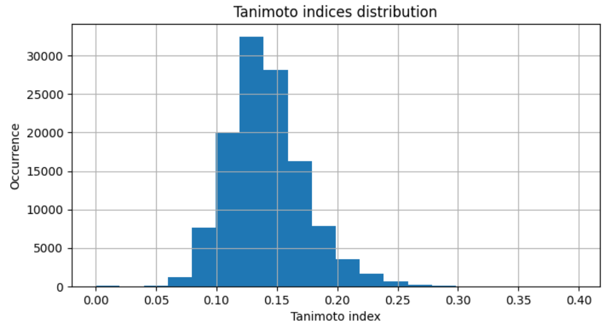 |
| CB1 | 1566 | CB1 | 314 | 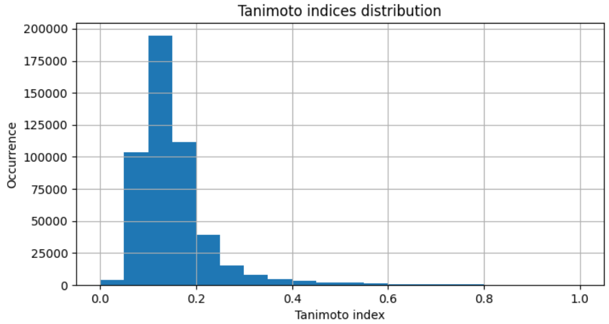 |
| | | CB2 | 418 | 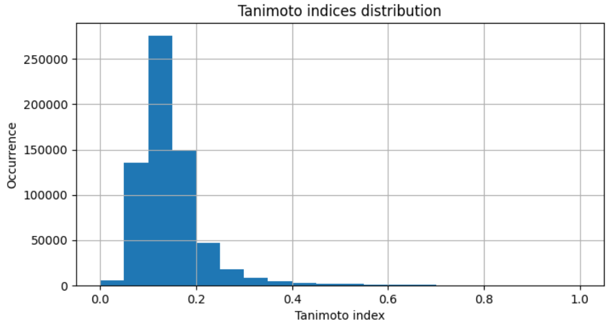 |
| | | CB2 selective | 35 | 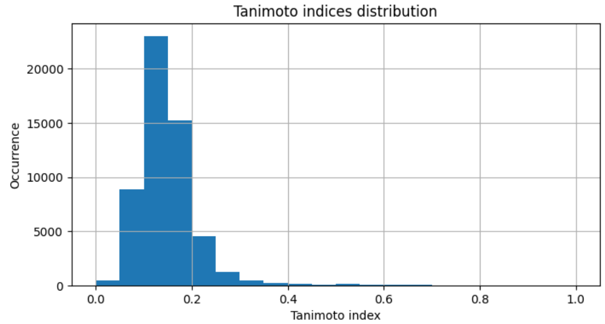 |

| | | | | |
|---|---|---|---|---|
| CB2 | 2093 | CB2 | 418 | 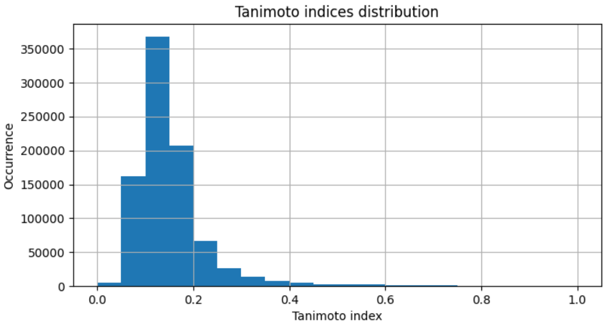  |
| | | CB1 | 314 | 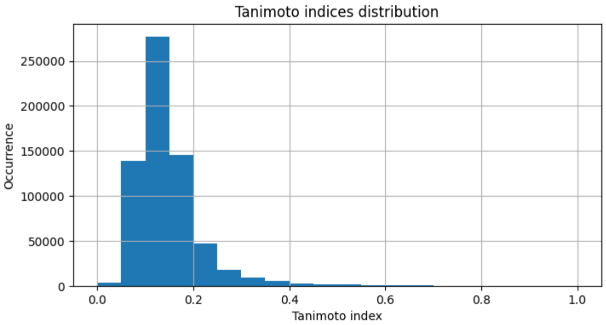  |
| | | CB2 selective | 35 | 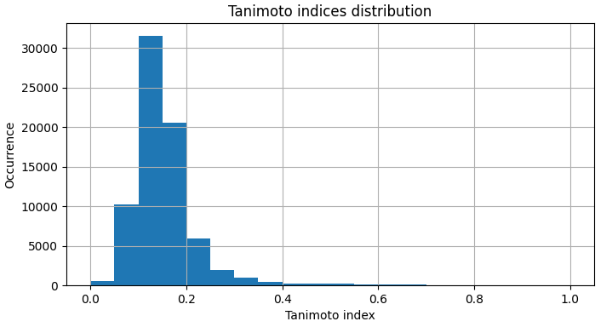  |

**Table S10.** Histograms of Tanimoto coefficients between the previous [4,5] and the current training datasets retrieved from ChEMBL.

| Previous training sets from https://db-gpcr.chem.uw.edu.pl | Number of datapoints | Current training sets including non-active compounds | Number of datapoints | Histogram of Tanimoto coefficients<br><br>Previous vs. current datasets |
|---|---|---|---|---|
| CB1 | 1566 | CB1 | 4509 | 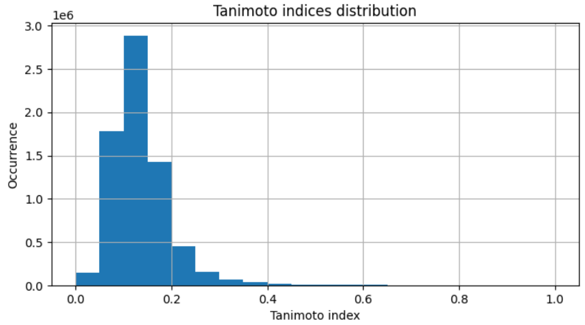  |
| CB2 | 2093 | CB2 | 4087 | 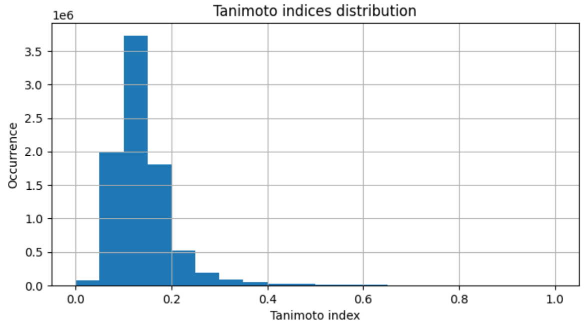  |
